# Serum glycoprotein markers in non-alcoholic steato-hepatitis and hepatocellular carcinoma

**DOI:** 10.1101/2021.09.30.462486

**Authors:** Prasanna Ramachandran, Gege Xu, Hector H. Huang, Rachel Rice, Bo Zhou, Klaus Lind-paintner, Daniel Serie

## Abstract

Fatty liver disease progresses through stages of fat accumulation and inflammation to non-alcoholic steatohepatitis (NASH), fibrosis and cirrhosis and eventually hepatocellular carcinoma (HCC). Currently available diagnostic tools for HCC lack sensitivity and specificity and deliver little value to patients. In this study, we investigated the use of circulating serum glycoproteins to identify a panel of potential prognostic markers that may be indicative of progression from the healthy state to NASH and further to HCC. Serum samples were processed using a standard pre-analytical sample preparation protocol and were analyzed using a novel high throughput glycoproteomics platform. We analyzed 413 glycopeptides, representing 57 abundant serum proteins and compared among the three phenotypes. Our initial dataset contained healthy, NASH, and HCC serum samples. We analyzed normalized abundance of common glycoforms and found 40 glycopeptides with statistically significant differences in abundances in NASH and HCC compared to controls. Summary level relative abundance of core-fucosylated, sialylated and branched glycans containing glycopeptides were higher in NASH and HCC as compared to controls. We replicated some of our findings in an independent set of samples of individuals with benign liver conditions and HCC, respectively. Our results may be of value in the management of liver diseases.

**TOC only:** 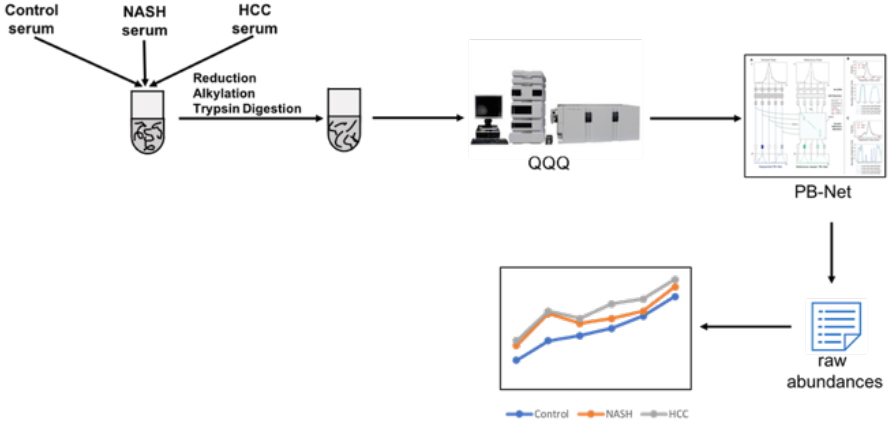

## Introduction

Accumulation of fat deposits in the liver, in the absence of excess alcohol consumption, is the hallmark of non-alcoholic fatty liver disease (NAFLD). NAFLD is the most common cause of chronic liver disease, affecting approximately 25% of the global population^1^. NAFLD progresses through various stages of fat accumulation from simple steatosis (NAFL) to steatosis and weak inflammation with or without fibrosis, a condition termed non-alcoholic steatohepatitis (NASH), which, in turn, may progress to the development of liver cirrhosis. Since about 1-2% of patients with liver cirrhosis will develop either end-stage liver diseases or hepatocellular carcinoma (HCC)^2–4^, early recognition of NAFLD and NASH represents an urgent unmet medical need. While liver biopsy is the gold standard and the most commonly used method for diagnosing NAFLD, its utility is limited by the invasive nature of the procedure as well as by the stochastic constraints imposed by histological heterogeneity^5, 6^.

A wide variety of noninvasive approaches have been developed for the noninvasive diagnosis of NAFLD and NASH, including imaging techniques, hepatic stiffness measurements using shear wave elastography or magnetic resonance elastography, and a multitude of biomarker-derived indices such as the aspartate aminotransferase-to-platelet ratio index (APRI), the FibroTest (gammaglutamyl transferase, total bilirubin, alpha-2-macroglobin (A2MG), apolipoprotein A1, and haptoglobin (HPT), with/without alanine aminotransferase [ALT]), the Firm index, the FibroIndex, the fibrosis-2 index, the Hui index, the NAFLD fibrosis score, or the BAAT-score (BMI, Age, ALT, triglycerides) ^7^). In addition, a large number of individual biomarkers including cytokeratin 18 (CK18)^8^, osteopontin^9^, fucosylated AFP (AFP-L3)^10^, des-gamma-gamma-carboxy prothrombin (DCP)^11^, glypican-3^12^, alpha-1-fucosidase^13^, Golgi protein-73 ^14^, alpha-1-acid-glycoprotein (AGP1)^15, 16^, alpha-fetoprotein (AFP)^17^, alpha-1-antitrypsin (A1AT) ^18, 19^, HPT ^18, 20–27^, apolipoprotein-J, A2MG, ceruloplasmin (CERU), CFAH, fibronectin, hemopexin (HEMO), kininogen, paraoxonase-1, vimentin, vitronectin (VTNC), mac-2-binding protein, immunoglobulin G (IgG)^28^, and miRNA^29^ have variably been cited as potentially useful to diagnose NAFLD/NASH and/or HCC; for the latter, AFP is used most widely^17^.

Common to all these indices and biomarkers is an underwhelming performance in real world testing, rendering them of limited utility and resulting in a multitude of missed diagnoses^30^. This is unfortunate, since NAFLD, and to a lesser extent NASH, in the absence of any approved pharmacologic treatments, may be reversible via simple dietary and lifestyle modifications if diagnosed early-on. Therefore, the development of an accurate, noninvasive diagnostic test for early recognition, with its expected major public health impact, has been the focus of numerous efforts.

Common to many of these putative biomarkers is that they are glycoproteins (cytokeratin 18, AGP1, AFP, A1AT, HPT, apolipoprotein-J, A2MG, CERU, CFAH, fibronectin, HEMO, kininogen, paraoxonase-1, vimentin, VTNC, mac-2-binding protein and IgGs). Indeed, higher levels of branching, sialylation and core fucosylation for a range of proteins have been found to be a hallmark of HCC^31^, and a “fucosylation index” has been considered as an indicator of progression from NASH to HCC^32^. Only a few detailed studies have been carried out investigating the association of shifts in relative abundance of individual glyco-isoforms of these proteins with the progression from the healthy state to NAFLD, NASH, and HCC. A recent publication by Zhu et al. found that characterization of HPT glycopeptide-isoforms might be useful in tracking progression from NASH/cirrhosis to early and late stage HCC^27^.

In this study, we applied a novel, high-throughput glycoproteomics platform to the interrogation of serum glycoprotein isoforms with the aim of finding clinically actionable, accurate biomarker panels that would allow for early, noninvasive recognition of, NAFLD/NASH as well as for monitoring the progression of fatty liver disorder to HCC.

## Materials and Methods

### Biological samples

The discovery set consisted of serum samples from 23 patients with a biopsy-proven diagnosis of NASH (10 male, 13 female; Indivumed AG, Hamburg, Germany) (Table1, Table S1), 20 patients with a diagnosis of HCC (16 male, 4 female; 6 stage I, 8 stage II, 6 stage III, 2 stage IV; Indivumed AG) (Table1, Table S2), and from 56 apparently healthy subjects with no history of liver disease (controls, 26 male, 30 female) which were sourced from iSpecimen (n=23, Lexington, MA), Palleon Pharmaceuticals Inc. (n=12, Waltham, MA) and Human Immune Monitoring Center (HIMC), Stanford University (n=21)) (Table1). Our validation set consisted of serum samples from 28 control subjects with a benign hepatic mass (16 male, 12 female) (Table1) and 28 subjects (20 male, 8 female) with HCC (Table 1), all obtained from Indivumed AG. Clinical diagnoses of patients with NASH and HCC were based on histopathological characterization of hepatic tissue obtained either via needle biopsy or at surgery.

**Table 1.** Summary of samples used in the discovery and validation sets

|  |  | Number of subjects | Male | Female | Age |
| --- | --- | --- | --- | --- | --- |
| Discovery | Control (healthy) | 56 | 26 | 30 | 23-91 |
|  | NASH | 23 | 10 | 13 | 45-70 |
|  | HCC | 19 | 15 | 4 | 32-85 |
| Validation | Control (benign hepatic mass) | 28 | 16 | 12 | 52-71 |
|  | HCC | 28 | 20 | 8 | 47-77 |

### Chemicals and reagents

Pooled human serum (for assay normalization and calibration purposes), dithiothreitol (DTT) and iodoacetamide (IAA) were purchased from Millipore Sigma (St. Louis, MO). Sequencing grade trypsin was purchased from Promega (Madison, WI). Acetonitrile (LC-MS grade) was purchased from Honeywell (Muskegon, MI). All other reagents used were procured from Millipore Sigma, VWR, and Fisher Scientific

### Preanalytical sample preparation

Serum samples were reduced with DTT and alkylated with IAA followed by digestion with trypsin in a water bath at 37°C for 18 hours. To quench the digestion, formic acid was added to each sample after incubation to a final concentration of 1% (v/v).

### Liquid chromatography/mass spectrometry (LC-MS) analysis

Digested serum samples were injected into an Agilent 6495B triple quadrupole mass spectrometer equipped with an Agilent 1290 Infinity ultra-high-pressure (UHP)-LC system and an Agilent ZORBAX Eclipse Plus C18 column (2.1 mm x 150 mm i.d., 1.8 μm particle size). Separation of the peptides and glycopeptides was performed using a 70-min binary gradient. The aqueous mobile phase A was 3% acetonitrile, 0.1% formic acid in water (v/v), and the organic mobile phase B was 90% acetonitrile 0.1% formic acid in water (v/v). The flow rate was set at 0.5 mL/min. Electrospray ionization (ESI) was used as the ionization source and was operated in positive ion mode. The triple quadrupole MS was operated in dynamic multiple reaction monitoring (dMRM) mode. Samples were injected in a randomized fashion with regard to underlying phenotype, and reference pooled serum digests were injected interspersed with study samples, at every 10^th^ sample position throughout the run.

### Data analysis

We performed MRM analysis of peptides and glycopeptides representing a total of 73 high-abundance serum glycoproteins. Our transition list consisted of glycopeptides as well as of non-glycosylated peptides from each glycoprotein. The python library Scikit-learn (https://scikit-learn.org/stable/) was used for all statistical analyses and for building machine learning models. We used PB-Net, a peak-integration software, that had been developed in-house to integrate peaks and to automatically obtain raw abundances for each marker^33^. Normalized abundance, corrected for within run drift, was calculated using the following formula:
Normalized abundance = (raw abundance of any glycopeptide or peptide in sample/raw abundance of a non-glycosylated peptide from the same glycoprotein) / average relative abundance of the same glycopeptides or peptides in the flanking pooled reference serum samples.

Relative abundance was calculated as the ratio of the raw abundance of any given glycopeptide to the sum of raw abundances of all glycopeptides.

Fold-changes for individual peptides and glycopeptides, were calculated on normalized abundances of control vs. NASH samples, control vs. HCC samples, as well as on NASH vs. HCC samples, after adjusting for age and sex. False discovery rate was calculated using the Benjamini-Hochberg method^34^. We performed principal component analysis (PCA) on normalized abundances of glycopeptides to investigate differences among the three phenotypes studied. Prior to performing PCA, normalized abundances were scaled so that the distribution had a mean value of 0 and a standard deviation of 1. Logistic regression models were built using normalized abundances of selected glycopeptides. The probability estimates of a sample in the test set predicted to belong to a particular phenotype was obtained from the trained logistic regression model.

### Ingenuity Pathway Analysis

Core analysis was performed to identify canonical pathways, up-stream regulators, and associated protein network by using Ingenuity^®^ Pathway Analysis (IPA) software (QIAGEN Inc.), relying on IPA’s proprietary algorithm to evaluate and minimize sample source bias. The p-value of overlap was calculated based on right-tailed Fisher’s exact test to determine the statistical significance of each canonical pathway, with p≤10^-3^ being considered statistically significant. The 10 statistically most significantly associated upstream regulators of differentially abundant glycoproteins identified in our study were predicted by using Ingenuity^®^ Knowledge Base. A molecule-class filter was applied to include only genes, RNAs and proteins. The networks associated with glycoproteins of interest were built based on both direct and indirect relationships. In addition, a total of 11 fucosyltransferase (FUT) genes and 20 sialyltransferase (ST) genes were retrieved from the CAZy database (www.cazy.org), and the IPA pathway explorer tool was used to explore the molecular connections of glycosylation-modifying enzymes and identified glycoproteins of interest. The “shortest path+1 node” was selected to construct the networks. Abundance values of the glycoproteins interrogated were not considered in these analyses.

## Results

### Normalized abundance of glycopeptides/peptides among control-, NASH-, and HCC-samples

We performed MRM analysis on control, NASH, and HCC serum samples. The peptide and glycopeptide markers employed in the MRM study were a selection of those published by Li et al^35^. The identity of each marker employed in our MRM experiments were verified by us. Figure S1 shows a representative example of chromatographic separation of different glycoforms of peptide VVLHPN*YSQVDIGLIK from HPT. In the MRM study of control, NASH, and HCC serum samples, normalized abundances of 187 glycopeptides and peptides were found to be statistically significantly different between samples from patients with NASH and controls with p-value of fold change less than 0.05. Likewise, normalized abundances of 254 glycopeptides and peptides were found to be statistically significantly different between samples from HCC patients and controls with p-values of fold change less than 0.05. Among these 254 glycopeptides and peptides, 215 showed differences that were statistically significant at a false discovery rate (FDR) of ≤0.05. Among the two sets of comparisons (NASH vs. controls, and HCC vs. controls), 87 glycopeptides and peptides were shared, i.e., showed statistically significantly different abundances in both comparisons at FDR<0.05. Among these 87 glycopeptides and peptides, the abundances of 40 glycopeptides and 23 peptides which exhibited statistically significantly differences that is also found in comparisons between samples from patients with NASH and controls. These 40 glycopeptides originated from 20 glycoproteins (Fig 1.a, Table S6). Likewise, normalized abundances of 166 glycopeptides and peptides were found to statistically significantly different between samples from NASH and HCC patients, with p-values of less than 0.05. Among these 72 glycopeptides and peptides, showed differences that were statistically significant at a false discovery rate FDR) of <0.05.

**Figure 1.**
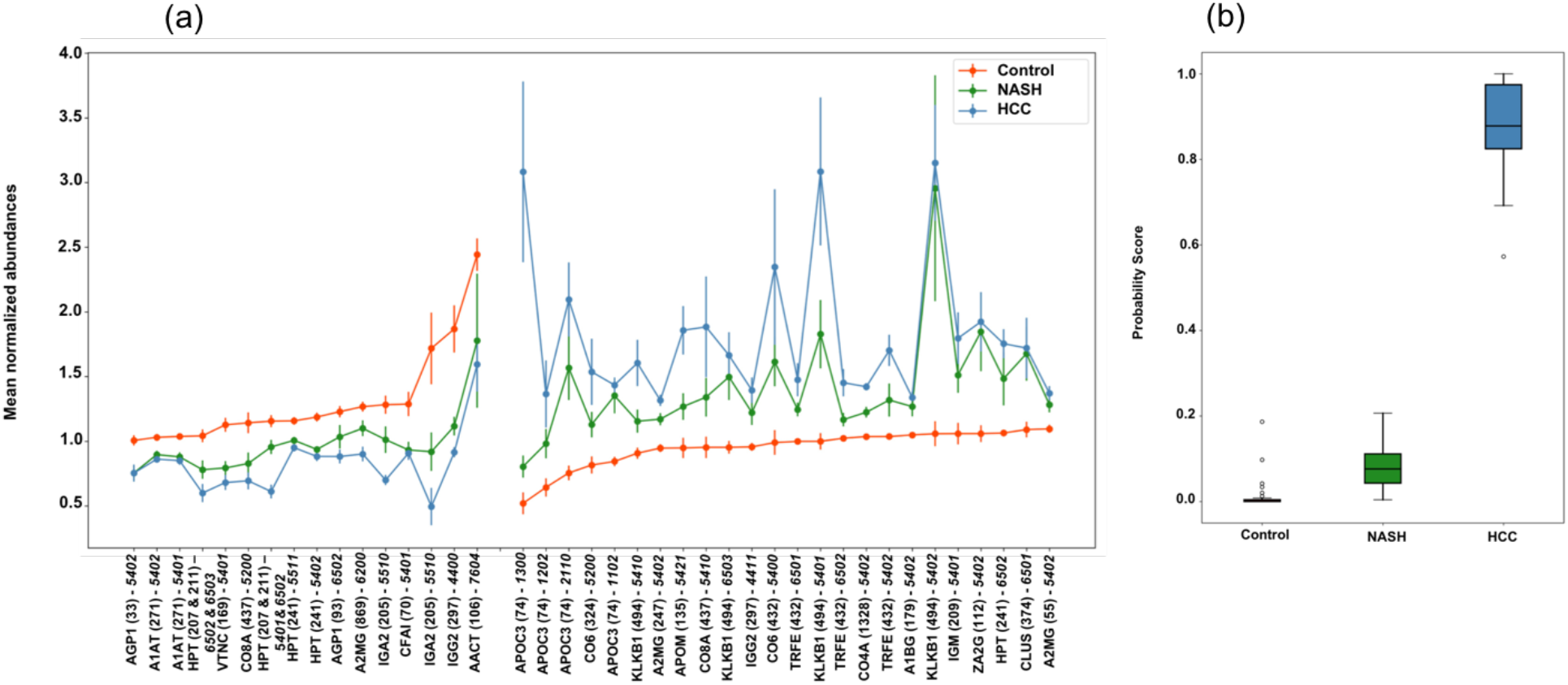
(a) Glycopeptide biomarkers in serum with progressive unidirectional changes in abundance of control-, NASH- and HCC-samples (b) Probability score for samples from control-, NASH- and HCC-subjects

Principal component analysis was performed to assess the segregation between the three phenotypes across first and second principal components (Fig. S2). While HCC samples segregate quite distinctly from control samples, most NASH samples do not. We trained a logistic regression model on normalized abundances of potential “disease progression markers”, i.e., glycopeptides/peptides that displayed unidirectionally higher or lower abundances across the phenotypic cascade from healthy to NASH to HCC. Fig 1.b shows the predicted probability of a sample representing the control, NASH, or HCC phenotype, respectively, based on this analysis. The coefficients of the logistic regression model are listed in Table S3. Among the 20 glycoproteins that were found to demonstrate statistically significant, unidirectional differences in abundance across the 3 phenotypes were seen in A2MG, HPT, apolipoprotein C3 (APOC3), CFAH, serotransferrin (TRFE), VTNC, CERU, A1AT. For differentiating glycoprotein profiles among NASH- and HCC-patients, we used logistic regression algorithm with LASSO regularization to build the model and leave one out cross validation (LOOCV) on NASH and HCC samples form the discovery set. We demonstrate an AUROC of 0.99 for the training set samples, and of 0.89 for the testing set (Figure S3).

### Relative abundance of glycopeptides containing common glycans among control-, NASH- and HCC-samples

We examined the cumulative relative abundances of glycopeptide motifs in control-, NASH- and HCC-samples. Higher levels of branching as well as of sialylation and core fucosylation have previously been reported for a range of proteins in HCC^31^. To further explore these findings, we examined glycopeptides with glycans containing no core fucosylation and either no sialylations (*0 Fuc, 0 Sial*), three sialylations (*0 Fuc, 3 Sial*), or four sialylations (*0 Fuc, 4 Sial*) among the glycopeptides identified as statistically significantly differentially abundant in our study. There were 49, 29, and 9 glycopeptides, respectively, in each of these 3 groups. We also examined glycopeptides with one core fucosylation and either two sialylations (*1 Fuc, 2 Sial*), three sialylations (*1 Fuc, 3 Sial*), or four sialylations (*1 Fuc, 4 Sial*) among the glycopeptides that are statistically significantly differentially abundant in our study (Fig 2). There were 33, 15, and 4 glycopeptides, respectively, in each of these 3 groups. Statistically significantly higher abundances were observed in relative abundance of all glycoforms with core fucosylation and multiple sialylations in NASH- and HCC-samples, respectively, as compared to control-samples. Statistically significant lower relative abundances of *0 Fuc, 3 Sial*-glycoforms were observed in NASH- and HCC-as compared to control-samples. Conversely, statistically significant higher abundances of *0 Fuc, 4 Sial*-glycoforms were observed in NASH- and HCC-samples as compared to control samples.

**Figure 2.**
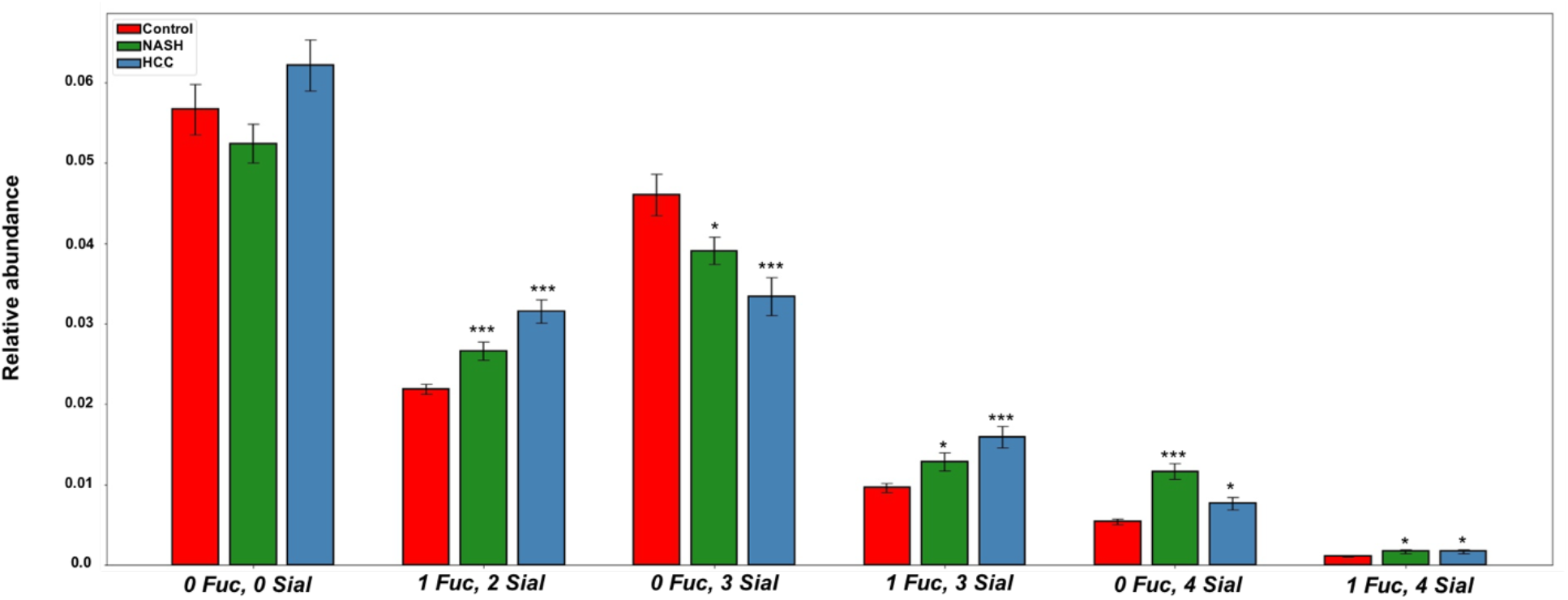
Relative abundances of common glycoforms by fucosylation and sialylation in control-, NASH- and HCC-samples. Columns indicate average relative abundances of glycans among the glycoproteins being monitored

Examination of the relative abundances of glycopeptides containing glycan moieties *5400, 5401, 5411* and *5412* revealed that abundances of those lacking core fucosylation (*5400* and *5401*) were statistically significantly less abundant in NASH- and HCC-samples as compared to control-samples. The abundances of glycans *5411* and *5412*, which contain core fucose and sialic acid residues, were statistically significantly more abundant in NASH- and HCC-samples as compared to control samples (Fig S4). We then analyzed the *65xx* series of glycoforms, which contain 5 N-acetyl-hexosamine (HexNaC), 6 hexose, and variable numbers of fucose and sialic acid residues, finding similar trends. Higher relative abundances were observed for sialylated and core-fucosylated glycopeptides, such as glycans *6511, 6512, 6513*, in HCC-samples as compared to control-samples. Statistically significantly higher relative abundances were observed for sialylated and core-fucosylated glycopeptides, such as glycans *6511*, *6513*, in NASH-samples as compared to control-samples. For glycoforms lacking core fucosylation but containing one or more sialylations, the result is more complex. Statistically significantly higher abundances were seen for *6501*, but statistically significant lower relative abundances were observed for *6502, 6503* in NASH- and HCC-samples as compared to control-samples (Fig S5). We also analyzed the *76xx* series of glycoforms that contain 6 HexNaC, 7 hexose and varying number of fucose and sialic acid residues. Relative abundances of multiply sialylated species *7602* and *7604* were statistically significantly much higher in NASH- and HCC-samples compared to control samples. Core fucosylated and multiply sialylated moieties *7613* and *7614* were statistically significantly more abundant in HCC-samples as compared to control-samples. Glycopeptides with glycan *7614* were statistically significantly more abundant in NASH compared to control samples. Meanwhile, their non-fucosylated, non-sialylated counterpart *7600* (Fig S6) showed no statistically significant difference among NASH- and HCC-samples as compared to control-samples.

### Glycoproteins with the most pronounced unidirectional quantitative differences among controls, NASH, and HCC

#### Alpha-2-macroglobulin (A2MG)

We observed statistically significant differences of four glycosylation sites (55, 247, 869 and 1424) for this protein (Fig 3, Table S4, Table S5). On site 1424, we found a statistically significantly lower abundance of glycan *5401* in HCC as compared to control samples. Glycan *5402*, containing no core fucosylation and two sialylations, was statistically significantly more abundant in NASH and HCC than in control patients at all 4 glycosylation sites. We observed statistically significantly lower abundances of the *5200*-glycan moiety at amino acid position 247 in HCC as compared to control samples. Likewise, glycans *5200*, *6200* and *6300* at amino acid position 869 displayed statistically significantly lower abundances in HCC as compared to controls. On the other hand, glycan *5401* was statistically significantly increased in HCC compared to control samples at site 869. Findings at amino acid position 55 were similar to those at amino acid position 1424 and 247. Glycan moiety *5402*, containing no core fucosylation and two sialylations, was statistically significantly more abundant in HCC-derived samples compared to samples derived from healthy subjects. At site 55, glycans *5411 and 5412* were statistically significantly less abundant in HCC cases as compared to controls. Also, statistically significantly higher abundances of A2MG protein were observed in in HCC patients as compared to controls (Fig 3, Table S4, Table S5).

**Figure 3.**
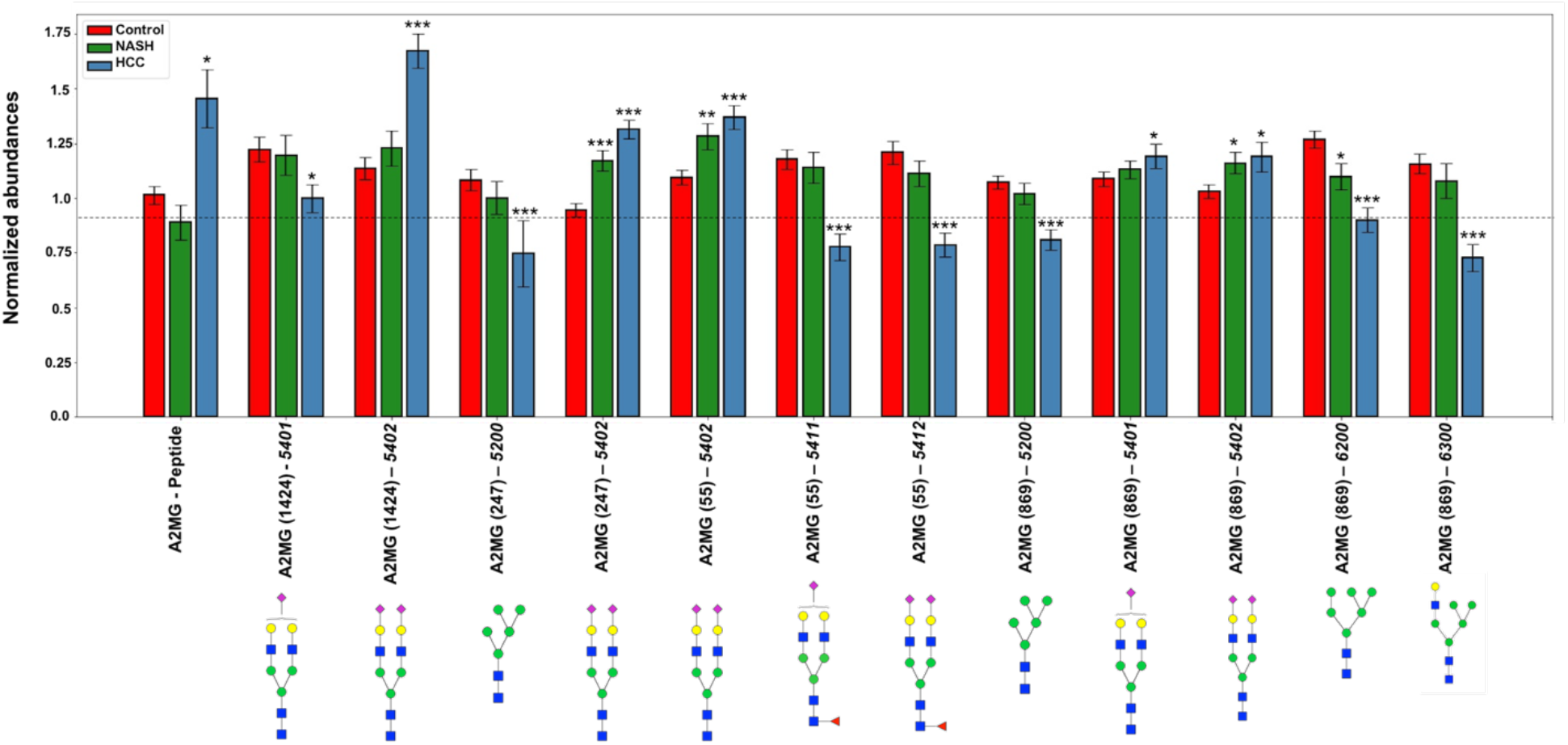
Normalized abundances of peptides and glycopeptides of A2MG in control-, NASH- and HCC-samples. Columns represent average normalized abundances of individual A2MG glycopeptides.

#### Alpha-1-acid glycoprotein 1 (AGP1)

The non-fucosylated, sialylated and tri-antennary (*6503*) glycopeptide at amino acid residue 103 was statistically significantly less abundant in HCC as compared to control samples (Fig S7, Table S4, Table S5). Meanwhile, the non-fucosylated, sialylated (*5402*) glycopeptide moiety at amino acid residue 33 was statistically significantly less abundant in NASH- and HCC-compared to control-samples. At amino acid site 93 statistically significantly lower abundances of moieties *6502* and *7604* (all lacking the core fucosylation) were observed in HCC as compared to control samples. Also, statistically significantly lower abundances of glycan moieties *6500* and *7604* were observed in in NASH-samples as compared to control samples on site 93. Moreover, statistically significantly higher abundances of glycans *7613* (containing a core fucose) were seen among HCC samples compared to controls at site 93. At amino acid residue 72, we observed statistically significantly lower abundances of glycan moiety *6503*, which lacks core fucosylation, in HCC as compared to control samples. At the same glycosylation site 72, statistically significantly higher abundances of branched, fucosylated, and multiply sialylated glycan moieties *7613, 7614* and *7601* (the latter lacking core fucosylation) were observed in HCC as compared to control samples (Fig S7, Table S4, Table S5).

#### Haptoglobin (HPT)

We evaluated at amino acid residue positions 184, 207 and 241 (Fig S8, Table S4, Table S5). At residue 184, we observed statistically significantly lower abundances of peptides carrying the non-fucosylated, mono-sialylated (*5401*) and mono-fucosylated, non-sialylated (*5410*) glycan motifs in HCC as compared to control. A statistically significantly higher abundance of glycans containing multiple sialic acid residues with (*5411*, *5412*) or without core-fucosylation (*5402*) and multiple sialylations were observed in HCC as compared to control samples. The peptide containing site 207 has multiple sites of glycosylation. The identity of individual glycans and site of attachment is not known. Out transition list also included a glycopeptide from haptoglobin with two sites of glycosylation. A statistically significant decrease in all glycoforms of the glycopeptide was observed in HCC compared to controls. A statistically significant decrease in three of these glycoforms was also observed in NASH compared to controls. At amino acid residue 241, statistically significantly lower abundances of glycan moieties *5401*, *5402*, *5511* were observed in NASH and HCC, as compared to control samples, while higher abundances of highly branched, sialylated, and core fucosylated glycan moieties (*6512, 6513, 7604*) were observed in HCC as compared to control samples (Fig S8, Table S4, Table S5).

#### Complement Factor H (CFAH)

At amino acid position 1029 we observed a statistically significantly lower abundance of glycan moieties *5401* and *5431* in HCC as compared to control samples. At site 882, we observed a statistically significantly lower abundances of glycans *5401* and *5402*, both of which lack core fucosylation, but are sialylated, in NASH and HCC as compared to control samples. Correspondingly, at this glycosylation site, a statistically significantly higher abundance of glycan *5411* was observed in HCC compared to control samples. At amino acid position 911, a statistically significantly higher abundance of doubly sialylated glycan moiety *5402*, along with a statistically significantly lower abundance of the singly sialylated glycan moiety *5401*, was observed in HCC as compared to control samples (Fig S9, Table S4, Table S5).

#### Alpha-1-antitrypsin (A1AT)

We observed statistically significantly higher abundances of core fucosylated, sialylated and branched glycans *6512* and *6513* at site 107 and of *5412* at site 271 and, correspondingly statistically significantly lower abundances of glycan species that lacked core fucosylation or sialylation, namely of 6502 at site 107 and of 5401 and 5402 at site 271, in NASH and HCC samples as compared to normal controls. Total levels of A1AT protein were statistically significantly increased in NASH compared to controls (Fig S10, Table S4, Table S5).

### Validation of results

We validated the results of the initial model by analyzing an independent set of samples from HCC patients and controls. The controls chosen were individuals with a diagnosis of a benign hepatic mass, to assess directly the discriminant power of differential glycopeptide abundance for HCC. In this set of samples, we were able to verify 12 glycopeptides and 2 of the peptides that had previously shown differences among healthy controls and HCC patients, with the directionality, magnitude of difference, and level of statistical significance being consistent among the 2 sample sets (Table 2, Fig 4). The 2 peptides and 9 of the 12 glycopeptides are associated with A2MG with the remaining 3 glycopeptides belonging to HPT, IGG1 and afamin (AFAM), respectively.

**Table 2.**
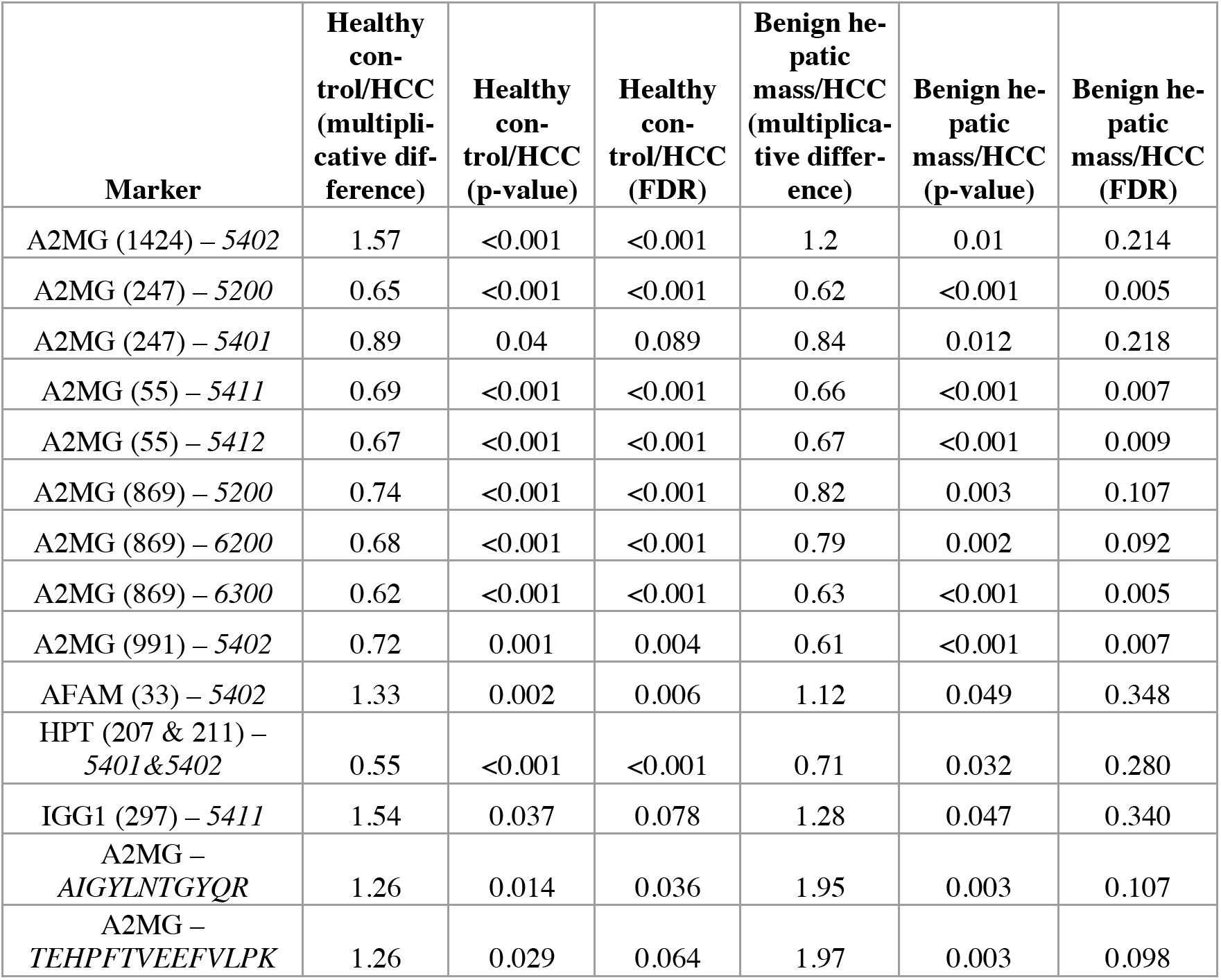
Glycopeptides displaying statistically significantly different abundances in both discovery and validation sample sets

| **Marker** | **Healthy control/HCC (multiplicative difference)** | **Healthy control/HCC (p-value)** | **Healthy control/HCC (FDR)** | **Benign hepatic mass/HCC (multiplicative difference)** | **Benign hepatic mass/HCC (p-value)** | **Benign hepatic mass/HCC (FDR)** |
| --- | --- | --- | --- | --- | --- | --- |
| A2MG (1424) – 5402 | 1.57 | <0.001 | <0.001 | 1.2 | 0.01 | 0.214 |
| A2MG (247) – 5200 | 0.65 | <0.001 | <0.001 | 0.62 | <0.001 | 0.005 |
| A2MG (247) – 5401 | 0.89 | 0.04 | 0.089 | 0.84 | 0.012 | 0.218 |
| A2MG (55) – 5411 | 0.69 | <0.001 | <0.001 | 0.66 | <0.001 | 0.007 |
| A2MG (55) – 5412 | 0.67 | <0.001 | <0.001 | 0.67 | <0.001 | 0.009 |
| A2MG (869) – 5200 | 0.74 | <0.001 | <0.001 | 0.82 | 0.003 | 0.107 |
| A2MG (869) – 6200 | 0.68 | <0.001 | <0.001 | 0.79 | 0.002 | 0.092 |
| A2MG (869) – 6300 | 0.62 | <0.001 | <0.001 | 0.63 | <0.001 | 0.005 |
| A2MG (991) – 5402 | 0.72 | 0.001 | 0.004 | 0.61 | <0.001 | 0.007 |
| AFAM (33) – 5402 | 1.33 | 0.002 | 0.006 | 1.12 | 0.049 | 0.348 |
| HPT (207 & 211) – 5401&5402 | 0.55 | <0.001 | <0.001 | 0.71 | 0.032 | 0.280 |
| IGG1 (297) – 5411 | 1.54 | 0.037 | 0.078 | 1.28 | 0.047 | 0.340 |
| A2MG – AIGYLNTGYQR | 1.26 | 0.014 | 0.036 | 1.95 | 0.003 | 0.107 |
| A2MG – TEHPFTVEEFVLPK | 1.26 | 0.029 | 0.064 | 1.97 | 0.003 | 0.098 |

**Figure 4.**
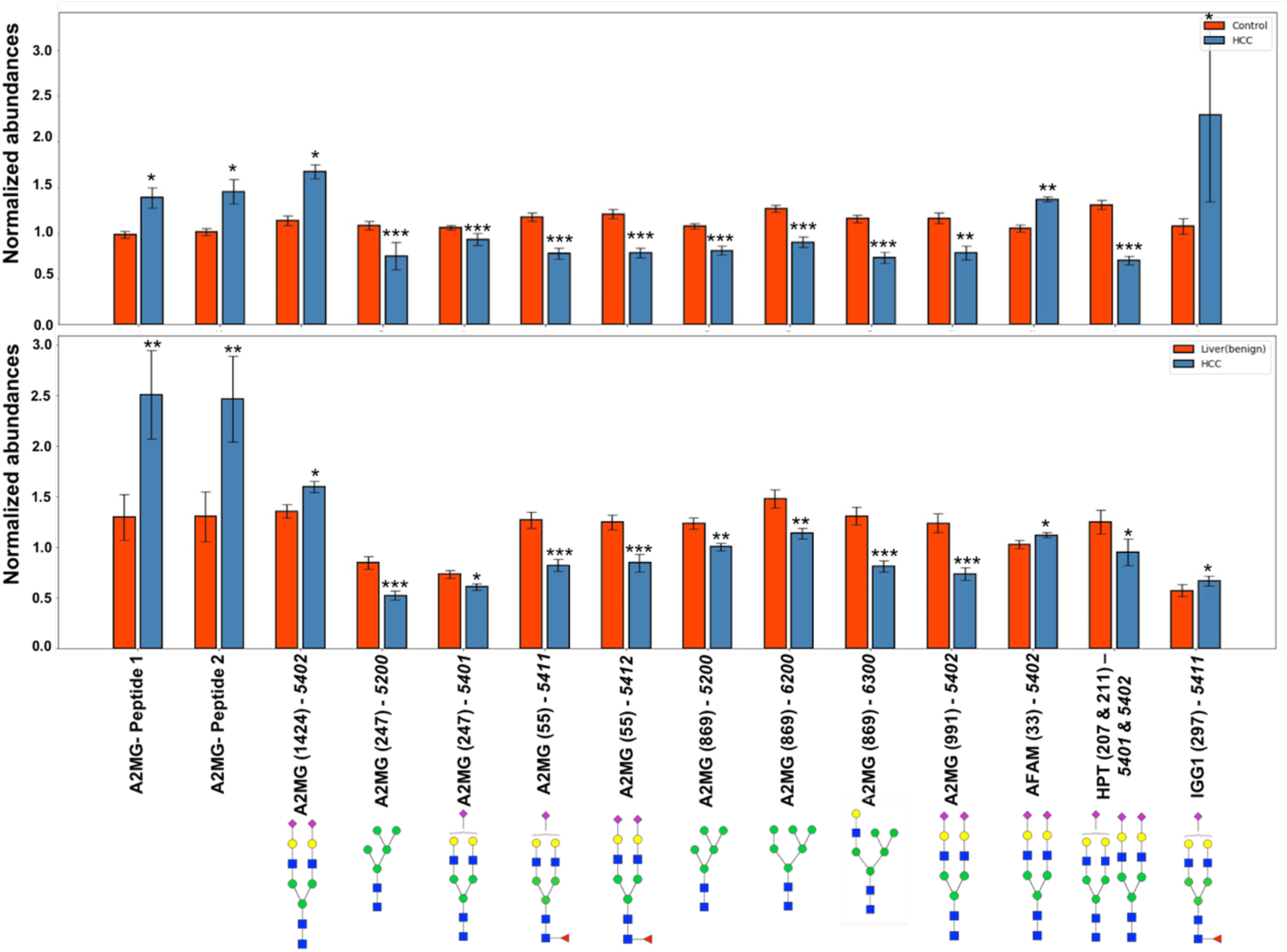
Normalized abundances of A2MG glycoforms in healthy controls and HCC, respectively, in discovery sample set (top panel). Normalized abundances of A2MG glycoforms in patients with benign hepatic masses and HCC, respectively, in validation sample set (bottom panel)

We built a logistic regression model using least absolute shrinkage and selection operator (LASSO) ^36^ regularization based on the samples of individuals with benign hepatic masses and of HCC patients, and performed a leave-one-out-cross-validation (LOOCV). We trained a LASSO model on all of the validation set except for one that left out, to test the model on. We tested the trained LASSO model on the data point that had been left out. We repeated this for every data point in the validation set. The consolidated results from LOOCV is represented in Fig 5. shows the receiver-operating-characteristic (ROC) curve for both the training and testing sets. The area under the ROC curve (AUROC) for the training set was found to be 0.85, and 0.77 for the testing set. When the LASSO model derived from the validation set was applied to the healthy controls and HCC samples from the discovery set, an AUROC of 0.87 was determined (Fig 5).

**Figure 5.**
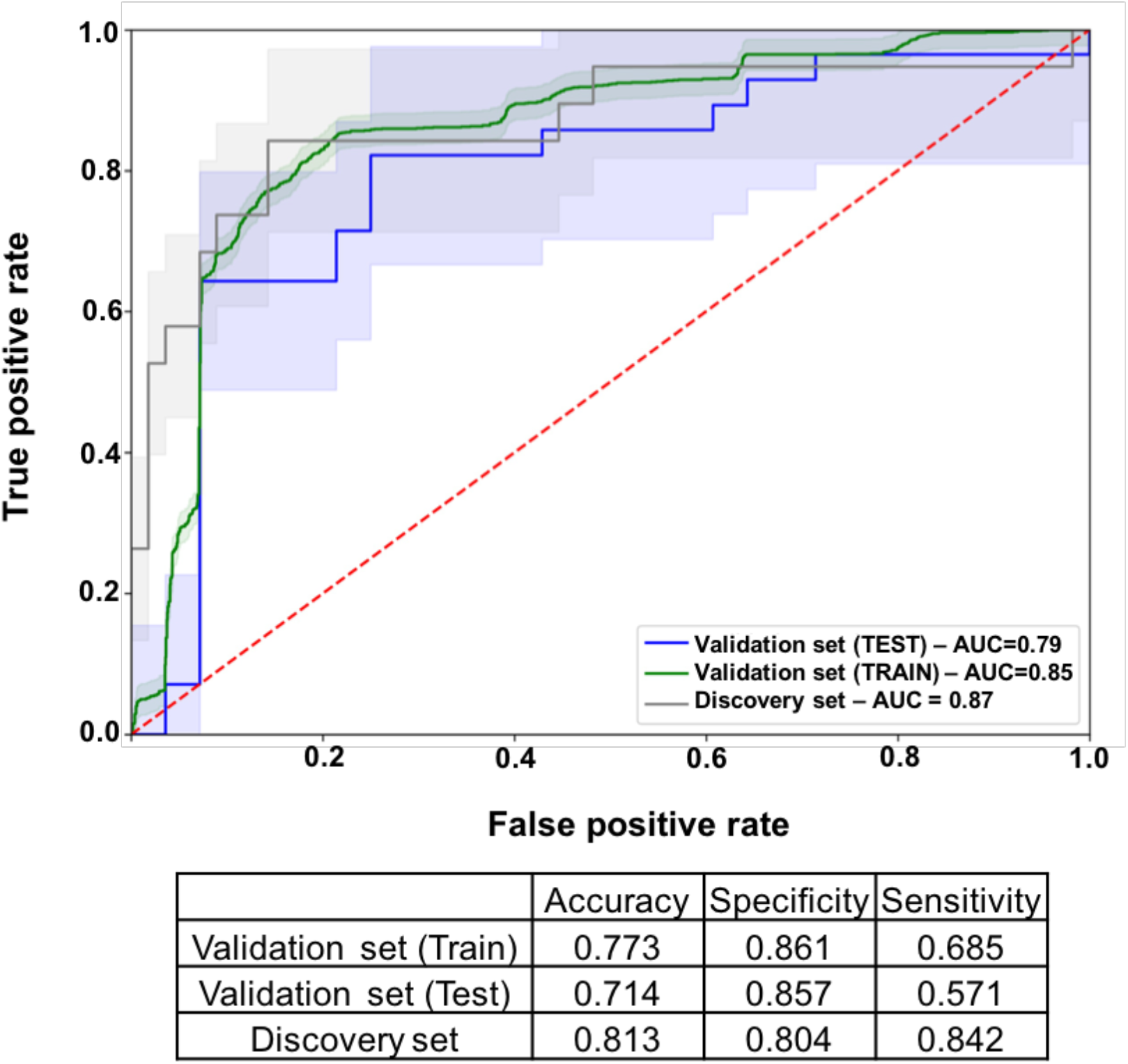
ROC curves generated using LOOCV in validation training and test sets, as well as applied to HCC and control samples in the discovery set

### Molecular pathway analysis

To explore functional biological aspects relevant for the 20 glycoproteins that were found to demonstrate statistically significant, unidirectional differences in glycopeptide abundance across the 3 phenotypes (Table S3), we performed IPA analysis to find canonical pathways, to discover potential regulatory networks, and to predict upstream regulators. The 10 statistically most significant canonical pathways with an overlapping p-value ≤10^-3^ are plotted in Fig 6.a, Table S6. The liver X receptor and retinoid acid X receptor (LXR/RXR) pathways, which are involved in regulating cholesterol and fatty acid metabolism, were identified as the most statistically significantly enriched pathways. Of the 20 glycoproteins interrogated, 9 are associated with this pathway, including A1BG, APOC3, CO4A/C4B, APOM, CLU, ORM1, SERPINA1, TF and VTNC. Additionally, the FXR/RXR pathway, acute phase response signaling, the complement system, and clathrin-mediated endocytosis signaling were among the 5 most enriched pathways. We next identified the 10 statistically most significantly associated upstream regulators for differentially abundant glycoproteins, using a p-value ≤10^-3^ as cutoff, including transcription regulators, transmembrane receptor, ligand dependent nuclear receptors and cytokines (Fig 6.b, Table S6). Solid lines in Fig 6.b represent a direct interaction between two molecules. Dotted lines represent an indirect interaction. Among the regulators thus identified, are hepatocyte nuclear factor 1α (HNF1α), hepatocyte nuclear factor 4α (HNF4α), and sterol regulatory element binding factor (SREBF1), three transcription factors prominently expressed in hepatocytes with multiple roles in the regulation of liver-specific genes. Dysregulation of HNF1α expression has been reported to be associated with both liver cirrhosis and hepatocellular carcinoma [34]. SREBF1 is involved in synthesis of cholesterol and lipids by regulating at least 30 pertinent genes^37^. The upstream regulator network, represented as a graph indicating the molecular relationships between these proteins, with the glycoproteins identified as statistically significantly abundant in our study highlighted in yellow (Fig 6.b). To gain further insight into molecular mechanisms associated with the N-linked glycosylation differences identified among these glycoproteins, 11 FUT and 20 ST genes were added to the analysis. The IPA Pathway explorer function was used to probe putative functional relationships of these glycosylation-modifying enzymes and the glycoproteins identified in our study as being of interest, based on the IPA Knowledge Base. Ten of the 11 FUT genes interrogated have been reported to be being directly or indirectly linked to glycoproteins identified in our study, via molecular intermediaries such as transcription factor HNF4α (Fig S11 (a)), and 12 of the 20 ST genes interrogated have been reported to affect 14 of the glycoproteins identified in our study, namely A2M, APOC3, AZGP1, C6, CFI, CLU, CO4A, IGHM, HP, ORM1, TF, SERPINA1, SERPINA3 and VTN via several transcription factors (e.g., SREBF1 and STAT6) or cytokines (e.g., IL1, IL2, IL6 and TNF) (Fig S11 (b)). These molecular networks indicate the potential crosstalk between several glycosyltransferases and the glycoproteins identified in our study.

**Figure 6.**
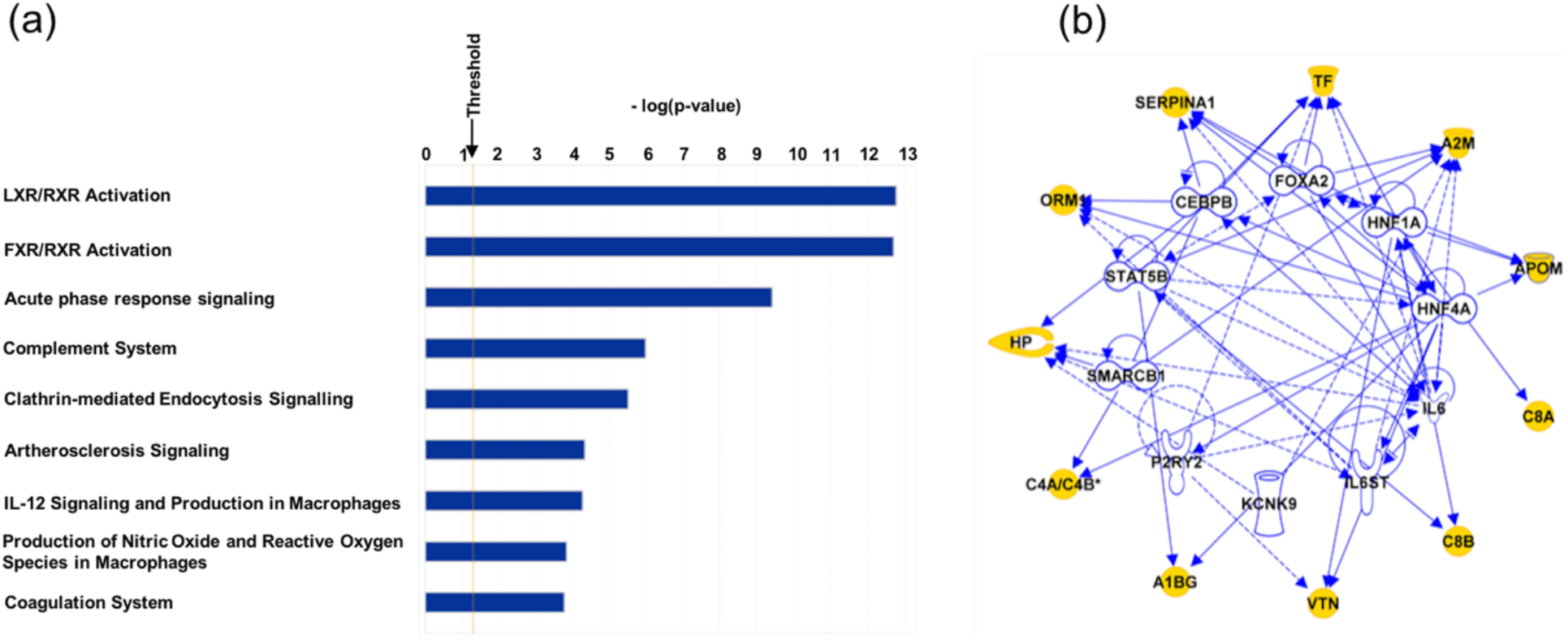
(a) Canonical pathways linked to proteins specified in Table 2 (IPA analysis). The horizontal bars represent the negative logarithm function of overlap p-value (b) Network of the 10 upstream regulator molecules statistically most significantly associated with genes encoding proteins specified in Table 2 (IPA analysis)

## Discussion

Our study is consistent with several previous studies that found higher relative abundance of core fucosylation, branching, and sialylation of glycans in NASH and HCC patients as compared to healthy controls. While many of the glycopeptides that we have identified as being associated with NASH had previously been reported in the literature, our study adds significant depth and detail for these biomarkers. These include APOC3 ^38^, apolipoprotein D (APOD) ^39^, apolipoprotein A1^40^, apolipoprotein M (APOM) ^41^, retinol binding protein-4^42^, HPT, A1AT, AGP1, VTNC, CFAH, IgA, IgG, IgM, hemopexin, TRFE ^43^, complement C8 alpha chain^44^ and A2MG^45^. Importantly, since a few of them (e.g., HPT ^27, 46^, A1AT ^47^, A2MG^48, 49^, and VTNC) have been reported previously as being differentially abundant at the protein level in NASH, our study opens important new insights into NASH biomarkers, as discussed below.

AGP1 has previously been studied as a potential biomarker for cirrhosis and HCC. Zhang et al. reported statistically significantly higher glycan branching, sialylation, and fucosylation of AGP1 glycopeptides in samples from patients suffering from NASH and cirrhosis as compared to controls ^15^. Several other studies have reported similar results for AGP1 glyco-isoforms in HCC ^16, 45, 50–52^. Our results confirm and expand these findings. We found higher normalized abundances of highly branched, core-fucosylated and multiply sialylated glycans in NASH and HCC as compared to healthy controls. Determination of the abundances of AGP-1 glycans may thus be of value when using this protein as a biomarker for NASH and HCC.

HPT has been proposed as a potentially useful marker for differentiating HCC from cirrhosis, with extensive work over the past few years highlighting, specifically, fucosylated haptoglobin as a marker for HCC and other liver diseases ^15, 20–24, 26, 27, 43, 53–55^. In all these studies, relatively higher levels of sialylated and fucosylated modifications of HPT in HCC as compared to controls have been reported. Moreover, HPT has also been evaluated as a marker for distinguishing NASH from hepatic steatosis ^56^. Kamada and coworkers found fucosylated and hyper-sialylated forms of HPT to be useful markers distinguishing NASH from NAFLD, and HCC from controls ^46, 56^. Our results confirm many of these findings and would justify further study of the use of HPT glyco-isoforms as markers for the diagnosis of NASH or HCC.

A1AT has previously been reported to be a marker for HCC. Communale et al. observed higher levels of glycans with core and outer arm fucosylation among 5 isoforms of A1AT^19^ in HCC as compared to healthy controls. Ahn et al. also reported higher levels of fucosylation of A1AT in HCC compared to hepatitis B virus (HBV) infected patients^57^. While decreased protein levels of A1AT in NAFLD compared to control healthy subjects have been reported in the past^47^, we found that A1AT protein levels were statistically significantly higher in NASH compared to controls.

APOC3 contains a single known O-glycosylation site. Overall protein levels of APOC3 have been reported to be lower in HCC^58^ compared to healthy controls. Our results are consistent with these findings. We found statistically significant lower levels of APOC3 protein in HCC compared to healthy controls. In addition, we found that levels were statistically significantly lower in NASH compared to healthy controls. We also found differences in O-glycosylation at amino acid position 74. While glycosylation variants of APOC3 have been reported to occur in breast cancer ^59^ and lung cancer ^60^, to our knowledge, our study is the first to demonstrate glycosylation differences of APOC3 in NASH and HCC.

CFAH has been extensively studied in HCC. Benicky and coworkers found that the ratios of fucosylated to non-fucosylated forms of the same glycan at amino acid residues 217, 882, 911 and 1029 ^61^ were higher in HCC as compared to controls. Darebna and coworkers observed higher core fucosylation levels at amino acid position 882^55^ in HCC as compared to controls, and our findings confirm these results. In addition, we found that the normalized abundance of core fucosylation is statistically significantly higher in NASH and in HCC, as compared to healthy controls. Contrary to a previous report^61^ based on a small number of samples and a different methodology, we found statistically significantly lower abundances of core-fucosylated glycopeptide species at amino acid residue 1029.

Specific glycopeptide moieties at amino acid position 1424 of A2MG have been reported to be present in the plasma of HCC patients^45^. We confirm this finding in our current study. Differential expression of A2MG glycoisoforms has also been reported in NASH patients^48, 49^. In our study, we demonstrate that A2MG glycoforms are associated with the progression from controls to NASH and to HCC and confirmed this trend in samples of patients with HCC compared to those with a benign hepatic mass. For several A2MG glycopeptides and peptides, the directionality and magnitude of differences across the spectrum from healthy controls to NASH and HCC appears representative of phenotype-aligned and phenotype-indicating progressive differences. We performed leave one out cross validation (LOOCV) on our validation set consisting of benign hepatic mass and HCC samples. Using logistic regression algorithm with LASSO regularization to build the model and LOOCV, we demonstrate an AUROC of 0.85 for the training set samples, and of 0.77 for the testing set. Subsequently, we built the LASSO model on the contrast of benign hepatic masses vs. HCCs using all samples in the validation set. When we used this trained model to predict on healthy controls vs. HCC, we determined an AUROC of 0.87, outperforming the validation set, test AUROC of 0.77 (Fig 5). This speaks to the robustness of glycopeptides as biomarkers distinguishing HCC from non-malignant liver conditions and from the healthy state.

Within the limitations inherent to the speculative nature of bioinformatics-based analyses, we highlight several plausible canonical pathways and upstream regulators linked to a selection of glycoproteins we found to have unidirectionally altered abundances among NASH and HCC samples. Likewise, we were able to demonstrate known interactions between a number of key enzymes involved in protein glycosylation and these glycoproteins. It is clear that these results, are at best suggestive of actual functional interactions and should be viewed as no more than hypothesis-generating; any more conclusive interpretation will have to await experimental confirmation.

The major shortcoming of the current study is the small sample size from patients with NASH that precluded splitting the cohort into a training and a testing set. Likewise, even though we were able confirm our findings with regard to HCC in an independent set of samples, the makeup of this second cohort (controls being individuals with benign hepatic lesions) was somewhat different from the first cohort (controls being healthy subjects without liver conditions). Additional work, using independent, ad ideally larger cohorts compatible with the phenotypes currently examined will be necessary to confirm our findings further. Another potential limitation of the present study is the fact that, based on the methods and protocol we applied, we are only interrogating a limited small number of relatively abundant serum glycoproteins; however, given the strength of our data, we believe that the advantage of a very simple workflow that lends itself to high throughput offsets the theoretical opportunity of obtaining even larger AuROCs.

## Conclusions

In summary, our work confirms previous findings demonstrating altered protein glycosylation in NASH and HCC. While previous studies explored either only single or few glycoproteins, we analyzed a large number of glycoproteins which resulted in the discovery of a broad panel of glycopeptide biomarkers associated with progression from the healthy state to NASH and ultimately HCC. This allowed us to build a highly accurate multivariable predictive classifier that clearly distinguishes between these conditions and that paves the way for generating a tool for early recognition of NASH and HCC. If confirmed in future prospective studies, our results may provide important new diagnostic tools in an area of currently unmet medical need.

## Supporting information

Supporting information in the manuscript

## Author Contributions

The manuscript was written through contributions of all authors. All authors have read and provided critical feedback, which shaped the research, analysis and manuscript and have given approval to the final version of the manuscript.

## Acknowledgements

We want to thank Palleon Pharmaceuticals Inc. and Human Immune Monitoring Center (HIMC) for providing healthy control samples for this study.

## Supporting information

**Figure S1.** Representative example for chromatographic separation of different glycoforms of the glycopeptide - VVLHPN*YSQVDIGLIK from haptoglobin

**Figure S2.** Principal component analysis of serum from control, NASH and HCC subjects using potential “progression markers”. The X-axis represents the first principal component and Y-axis represents the second principal component. Each dot represents first and second principal component coordinates of a subject.

**Figure S3.** ROC curve from leave-one-out cross validation on NASH and HCC samples in the discovery dataset.

**Figure S4.** Relative abundance of common glycoforms *5400, 5401, 5411, 5412* in control, NASH and HCC serum across all 73 glycoproteins studied. Columns indicate cumulative relative abundances of glycans among the glycoproteins being monitored.

**Figure S5.** Relative abundances of common glycoforms *6501, 6511, 6512, 6502, 6512, 6503, 6513* in control, NASH, and HCC serum across all 73 glycoproteins studies. Columns indicate cumulative relative abundances of glycans among the glycoproteins being monitored.

**Figure S6.** Relative abundances of common glycoforms *7600, 7602, 7604, 7613, 7614* in control, NASH and HCC serum across all 73 glycoproteins studied. Columns indicate cumulative relative abundances of glycans among the glycoproteins being monitored.

**Figure S7.** Normalized abundances of peptide and glycopeptides of AGP1 in control, NASH, and HCC serum across all 73 glycoproteins studied. Columns indicate normalized abundances of a certain type of glycans.

**Figure S8.** Normalized abundances of peptide and glycopeptides of HPT in control-, NASH- and HCC-samples. Columns indicate normalized abundances of a certain type of glycans.

**Figure S9.** Normalized abundances of peptide and glycopeptides of CFAH in control-, NASH- and HCC-samples. Columns indicate normalized abundances of glycans.

**Figure S10.** Normalized abundance of peptide and glycopeptides of A1AT in control-, NASH- and HCC-samples. Columns indicate normalized abundances of glycans.

**Figure S11.** (a) Network of fucosyltransferases and target glycoproteins. Solid lines represent a direct interaction between molecules. Dotted lines represent an indirect interaction. (b) Network of sialyltransferases and target glycoproteins

**Table S1.** Summary of NASH patients in the discovery dataset

**Table S2.** Summary of HCC patients in the discovery dataset

**Table S3.** Multiplicative differences, Student’s t-test p-values, and FDR values for unidirectionally differentially expressed glycopeptides (“progression markers”)

**Table S4.** Multiplicative difference between NASH/control and HCC/control

**Table S5.** Glycan code and structure

**Table S6.** IPA analysis - top ten upstream regulators

