## Supporting information in the manuscript for "Serum glycoprotein markers in non-alcoholic steato-hepatitis and hepatocellular carcinoma"

**Figure S1.** Representative example for chromatographic separation of different glycoforms of the glycopeptide - VVLHPN*YSQVDIGLIK from haptoglobin

**Figure S2.** Principal component analysis of serum from control, NASH and HCC subjects using potential "progression markers". The X-axis represents the first principal component and Y-axis represents the second principal component. Each dot represents first and second principal component coordinates of a subject.

**Figure S3.** ROC curve from leave-one-out cross validation on NASH and HCC samples in the discovery dataset.

**Figure S4.** Relative abundance of common glycoforms *5400, 5401*, *5411*, *5412* in control, NASH and HCC serum across all 73 glycoproteins studied. Columns indicate cumulative relative abundances of glycans among the glycoproteins being monitored.

**Figure S5.** Relative abundances of common glycoforms *6501*, *6511*, *6512*, *6502*, *6512*, *6503*, *6513* in control, NASH, and HCC serum across all 73 glycoproteins studies. Columns indicate cumulative relative abundances of glycans among the glycoproteins being monitored.

**Figure S6.** Relative abundances of common glycoforms *7600*, *7602, 7604*, *7613*, *7614* in control, NASH and HCC serum across all 73 glycoproteins studied. Columns indicate cumulative relative abundances of glycans among the glycoproteins being monitored.

**Figure S7.** Normalized abundances of peptide and glycopeptides of AGP1 in control, NASH, and HCC serum across all 73 glycoproteins studied. Columns indicate normalized abundances of a certain type of glycans.

**Figure S8.** Normalized abundances of peptide and glycopeptides of HPT in control-, NASH- and HCC-samples. Columns indicate normalized abundances of a certain type of glycans.

**Figure S9.** Normalized abundances of peptide and glycopeptides of CFAH in control-, NASH- and HCC-samples. Columns indicate normalized abundances of glycans.

**Figure S10.** Normalized abundance of peptide and glycopeptides of A1AT in control-, NASH- and HCC-samples. Columns indicate normalized abundances of glycans.

**Figure S11.** (a) Network of fucosyltransferases and target glycoproteins. Solid lines represent a direct interaction between molecules. Dotted lines represent an indirect interaction. (b) Network of sialyltransferases and target glycoproteins

**Table S1.** Summary of NASH patients in the discovery dataset

**Table S2.** Summary of HCC patients in the discovery dataset

**Table S3.** Multiplicative differences, Student's t-test p-values, and FDR values for unidirectionally differentially expressed glycopeptides ("progression markers")

**Table S4.** Multiplicative difference between NASH/control and HCC/control

**Table S5.** Glycan code and structure

**Table S6.** IPA analysis - top ten upstream regulators

**Fig S1.**

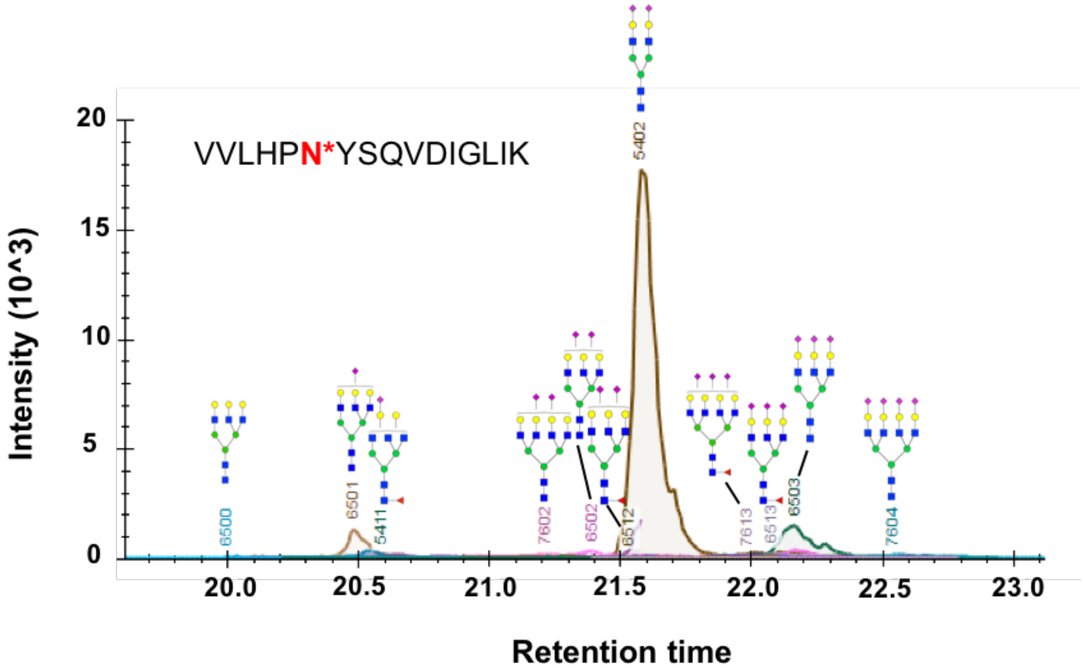
VVLHPN*YSQVDIGLIK
Intensity (10^3)
20
15
10
5
0
Retention time
20.0
20.5
21.0
21.5
22.0
22.5
23.0
6500
6501
5411
7602
6502
6512
5402
7613
6513
6503
7604

**Figure S2.**

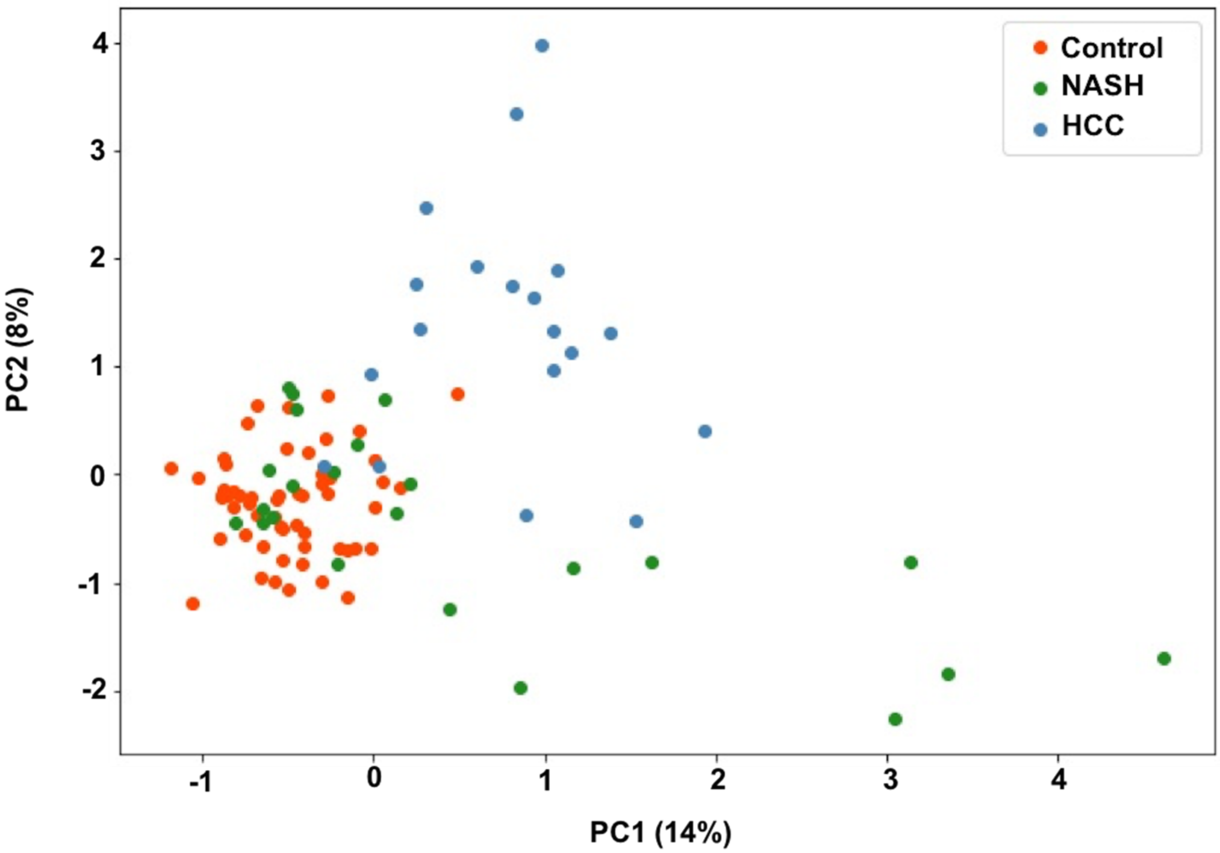

**Figure S3**

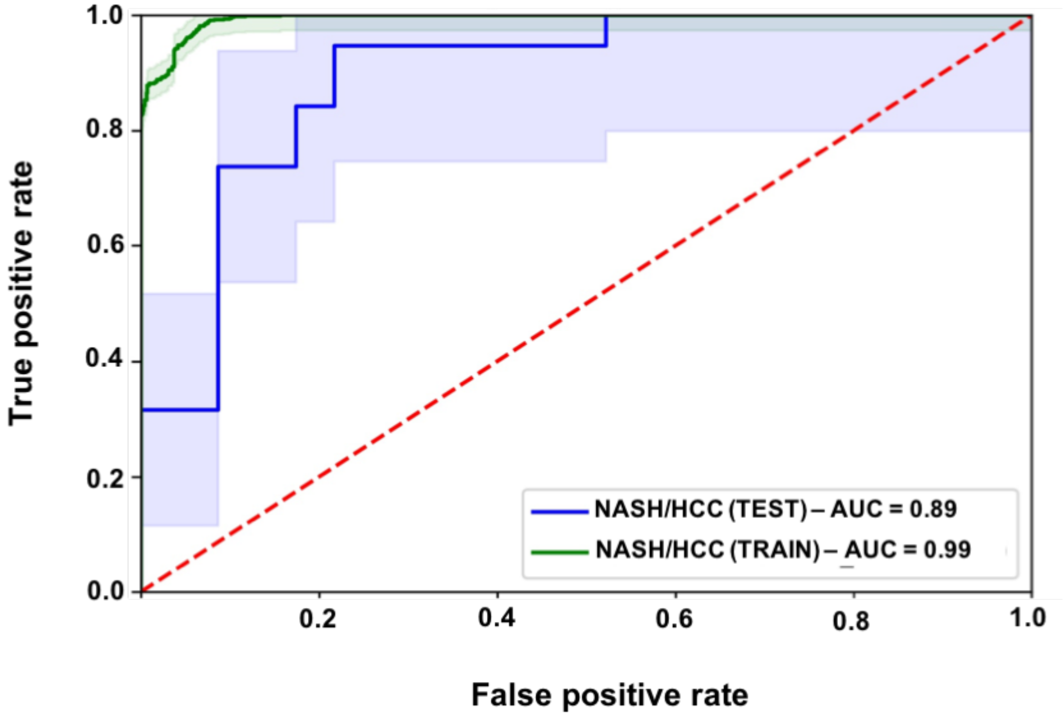


| | Accuracy | Specificity | Sensitivity |
|---|---|---|---|
| NASH/HCC (Train) | 0.952 | 0.861 | 0.949 |
| NASH/HCC (Test) | 0.857 | 0.783 | 0.947 |

**Figure S4.**

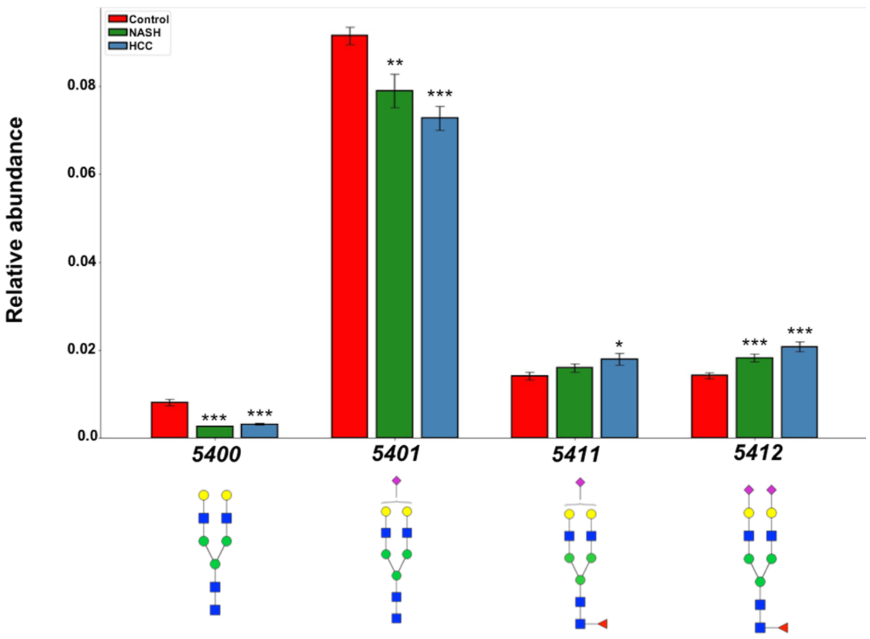


**Figure S5.**

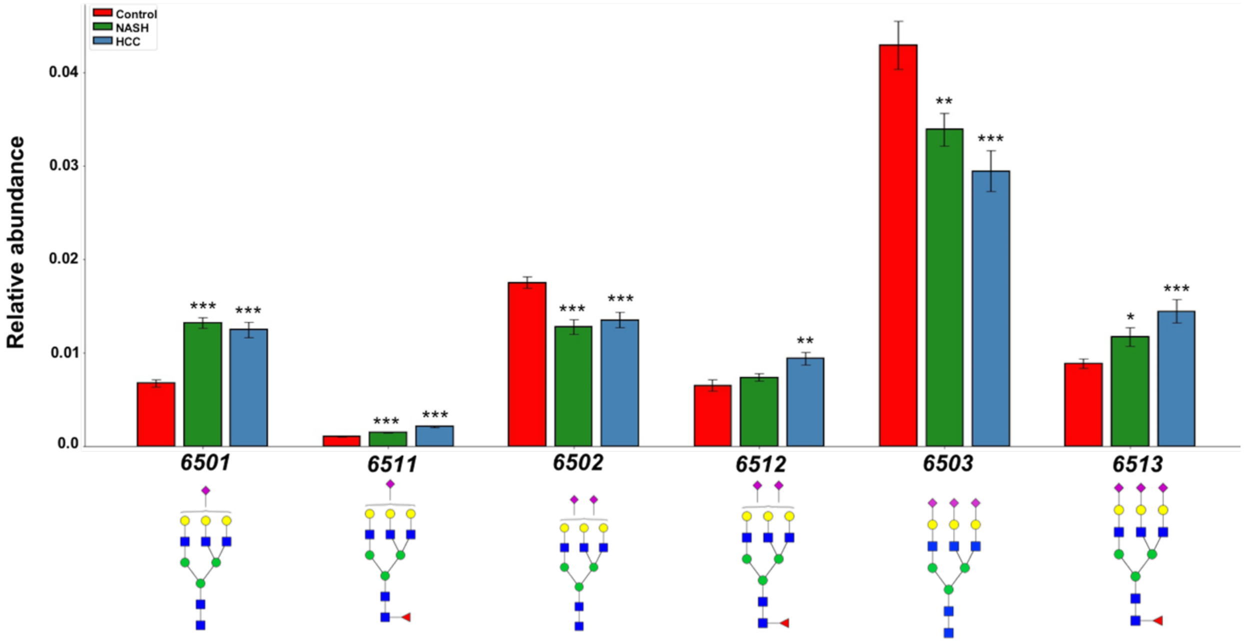

**Figure S6.**

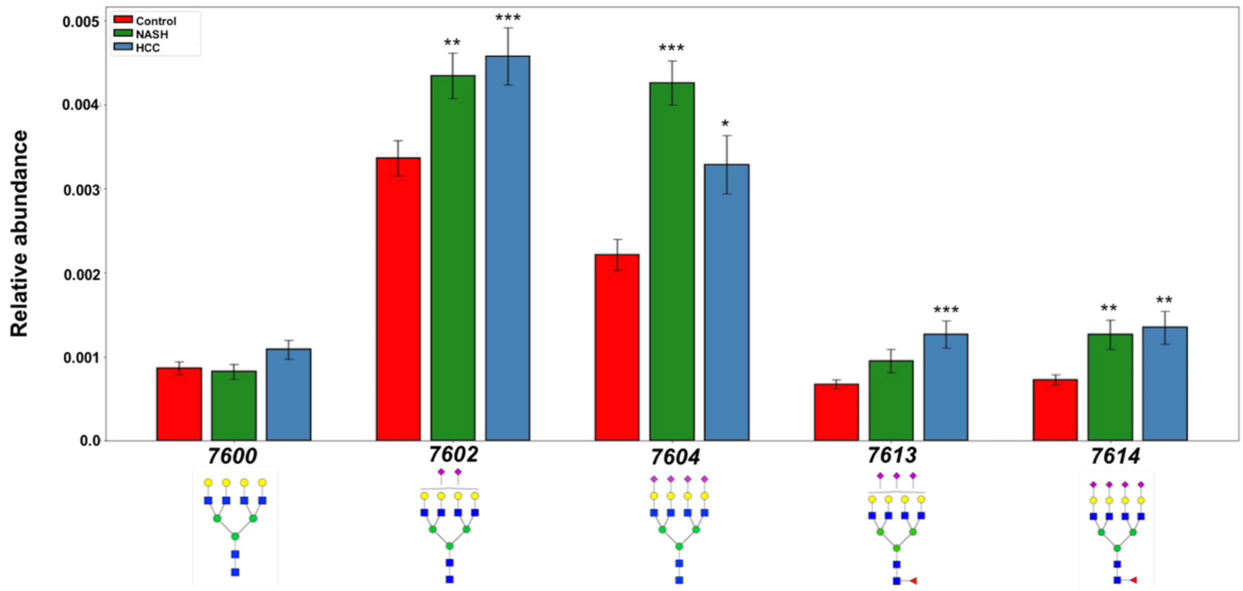


**Figure S7.**

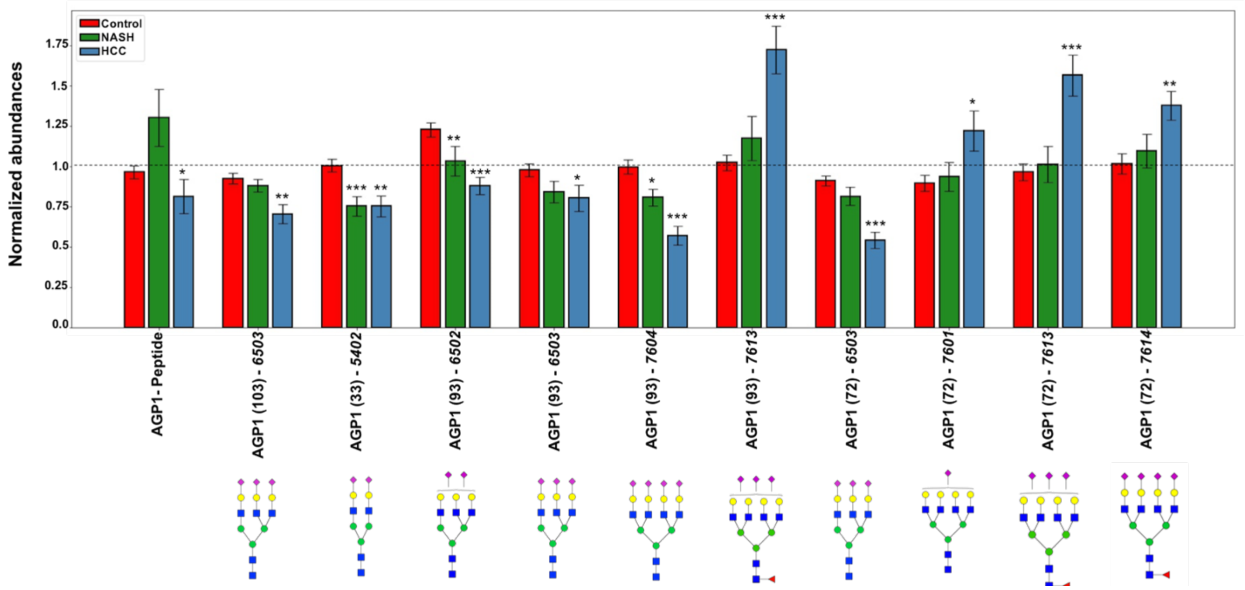

### Figure S8.

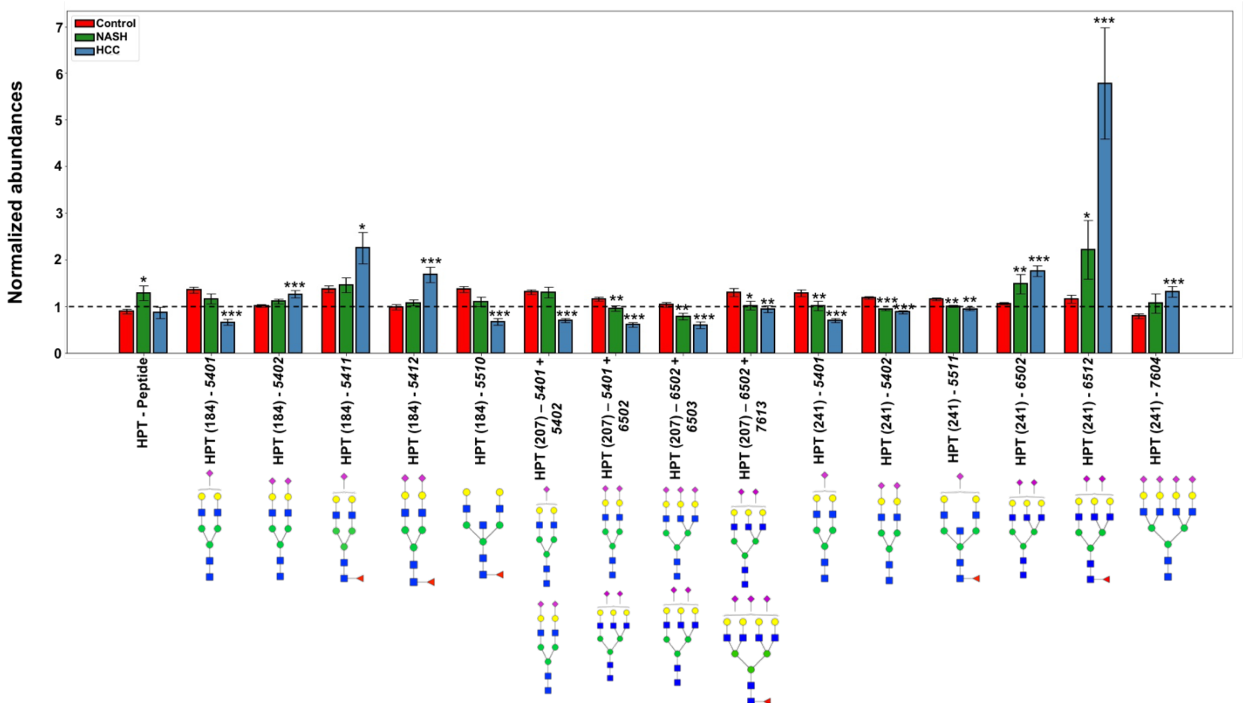


### Figure S9.

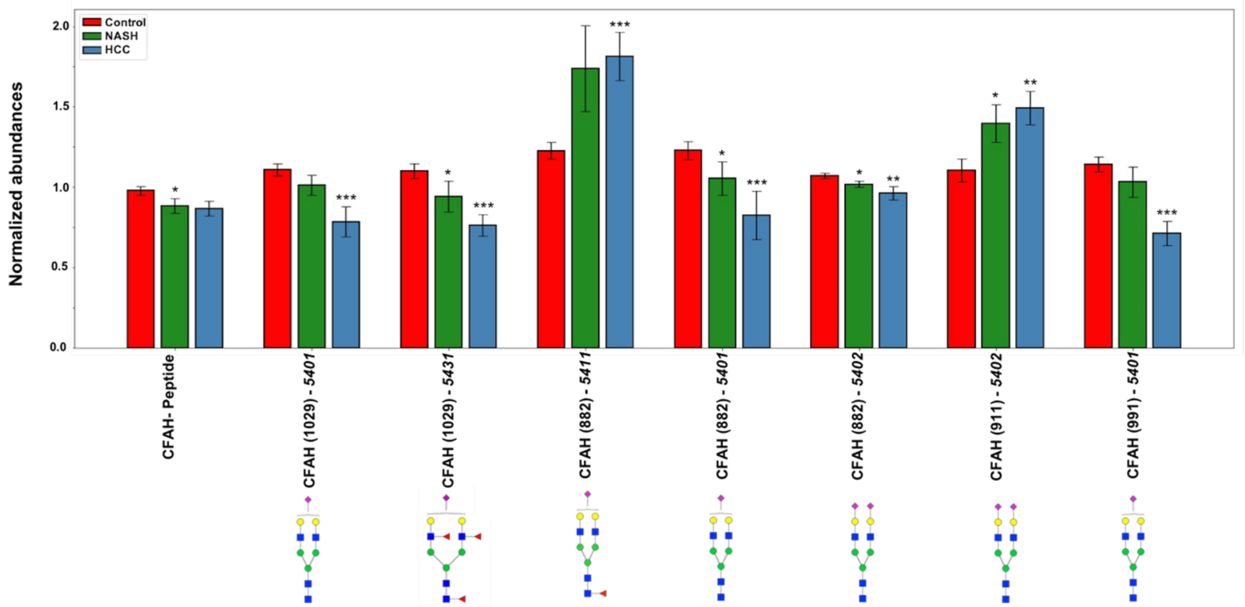

**Figure S10.**

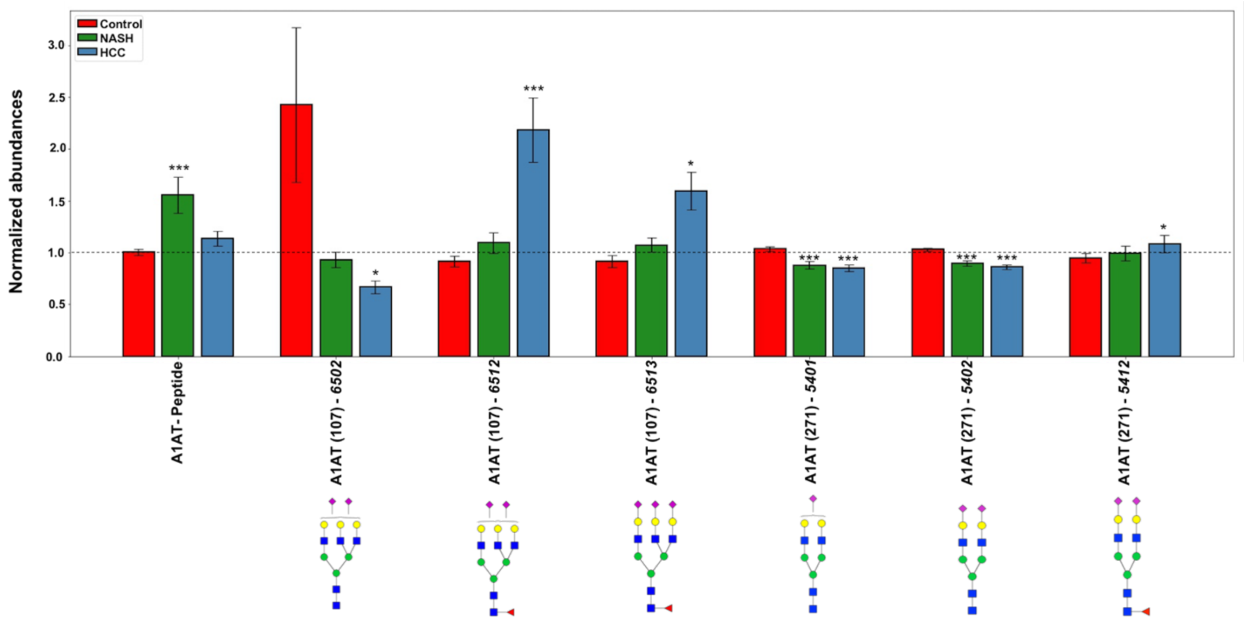


**Figure S11.**

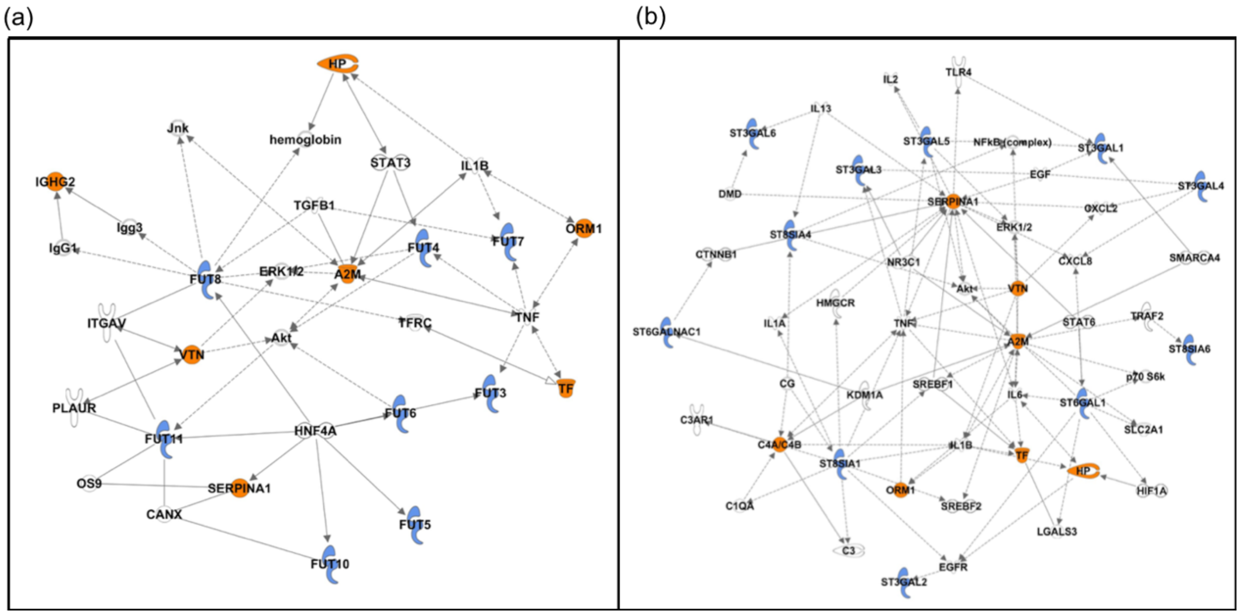

### Table S1.

| | Number of females | Number of males |
|---|---|---|
| **Fibrosis stage** | | |
| 1B | 13 | 10 |
| **Steatosis** | | |
| 2 | 10 | 7 |
| 3 | 3 | 3 |
| **Lobular inflammation** | | |
| 2 | 9 | 7 |
| 3 | 4 | 3 |
| **Ballooning** | | |
| 1 | 1 | |
| 2 | 4 | 6 |
| 3 | 8 | 4 |

### Table S2.

| | Number of females | Number of males |
|---|---|---|
| **Tumor size (cm)** | | |
| < 5 cm | 1 | 5 |
| >5 cm | 2 | 9 |
| missing | 1 | 2 |
| **T_classification** | | |
| T1 | 1 | 6 |
| T2 | 2 | 6 |
| T3 | 1 | 4 |
| **N_classification** | | |
| N0 | 3 | 15 |
| N1 | 1 | 1 |
| **M_classification** | | |
| M0 | 4 | 16 |
| **Stage** | 4 | 6 |
| 1 | | 6 |
| II | 2 | 6 |
| III | 1 | 3 |
| IV | 1 | 1 |

| Grading | | |
| --- | --- | --- |
| G1 | 1 | 2 |
| G2 | 1 | 6 |
| G3 | 2 | 5 |
| G4 | | 1 |
| missing | | 2 |

### Table S3.

| Marker | protein | gene | control/ NASH (multiplicative difference) | control /NASH (p-value) | control/ NASH (FDR) | control/ HCC (multiplicative difference) | control /HCC (p-value) | control /HCC (FDR) | Logistic regression model coefficients (NASH vs Rest) | Logistic regression model coefficients (HCC vs Rest) |
| --- | --- | --- | --- | --- | --- | --- | --- | --- | --- | --- |
| A1AT (271) - *5401* | A1AT | SERPINA1A | 0.85 | <0.001 | 0.005 | 0.79 | <0.001 | <0.001 | 0.013 | -0.412 |
| A1AT (271) - *5402* | A1AT | SERPINA1A | 0.87 | <0.001 | <0.001 | 0.82 | <0.001 | <0.001 | 0.099 | -0.369 |
| A1BG (179) - *5402* | A1BG | A1BG | 1.17 | <0.001 | 0.003 | 1.28 | <0.001 | <0.001 | 0.17 | 0.056 |
| A2MG (247) - *5402* | A2MG | A2M | 1.23 | <0.001 | 0.002 | 1.47 | <0.001 | <0.001 | -0.045 | 0.235 |
| A2MG (55) - *5402* | A2MG | A2M | 1.17 | 0.003 | 0.016 | 1.34 | <0.001 | <0.001 | -0.108 | 0.327 |
| A2MG (869) - *6200* | A2MG | A2M | 0.87 | 0.013 | 0.049 | 0.68 | <0.001 | <0.001 | 0.576 | -0.435 |
| AACT (106) - *7604* | AACT | SERPINA3 | 0.45 | <0.001 | 0.001 | 0.63 | 0.003 | 0.009 | -0.287 | -0.0507 |
| AGP1 (33) - *5402* | AGP1 | ORM1 | 0.72 | <0.001 | 0.003 | 0.75 | 0.002 | 0.008 | -0.138 | 0.11 |
| AGP1 (93) - *6502* | AGP1 | ORM1 | 0.82 | 0.009 | 0.038 | 0.69 | <0.001 | <0.001 | 0.119 | -0.701 |
| APOC3 (74) - *1102* | APOC3 | APOC3 | 1.53 | <0.001 | <0.001 | 1.77 | <0.001 | <0.001 | 0.312 | -0.123 |
| APOC3 (74) - *1202* | APOC3 | APOC3 | 1.73 | 0.001 | 0.008 | 2.55 | <0.001 | <0.001 | 0.31 | 0.071 |
| APOC3 (74) - *1300* | APOC3 | APOC3 | 2.01 | <0.001 | 0.005 | 7.53 | <0.001 | <0.001 | -1.44 | 0.588 |

| | | | | | | | | | | |
|---|---|---|---|---|---|---|---|---|---|---|
| APOC3 (74) - *2110* | APOC3 | APOC3 | 1.93 | <0.001 | <0.001 | 2.98 | <0.001 | <0.001 | 0.55 | 0.391 |
| APOM (135) - *5421* | APOM | APOM | 1.62 | 0.014 | 0.049 | 2.3 | <0.001 | 0.002 | -0.14 | 0.624 |
| CFAI (70) - *5401* | CFAI | CFI | 0.76 | 0.004 | 0.024 | 0.74 | 0.004 | 0.0124 | -0.68 | 0.075 |
| CLUS (374) - *6501* | CLUS | CLU | 1.46 | <0.001 | 0.005 | 1.53 | <0.001 | 0.003 | 1.119 | -0.644 |
| CO4A (1328) – *5402* | CO4A | C4A | 1.17 | <0.001 | <0.001 | 1.39 | <0.001 | <0.001 | 0.398 | 0.45 |
| CO6 (324) - *5200* | CO6 | C6 | 1.57 | 0.005 | 0.025 | 2.42 | <0.001 | <0.001 | -0.295 | 0.082 |
| CO6 (324) - *5400* | CO6 | C6 | 1.76 | 0.002 | 0.01 | 1.9 | 0.008 | 0.023 | 0.186 | 0.101 |
| CO8A (437) - *5200* | CO8A | C8A | 0.72 | 0.011 | 0.043 | 0.57 | <0.001 | <0.001 | -0.277 | -0.974 |
| CO8A (437) - *5410* | CO8A | C8A | 1.43 | 0.008 | 0.035 | 1.75 | 0.002 | 0.006 | 0.274 | -0.122 |
| HPT (207) - *5401+ 6502* | HPT | HP | 0.82 | 0.006 | 0.031 | 0.54 | <0.001 | <0.001 | 0.769 | -1.036 |
| HPT (207) - *6502+6503* | HPT | HP | 0.72 | 0.005 | 0.025 | 0.56 | <0.001 | <0.001 | 0.335 | -0.606 |
| HPT (241) - *5401* | HPT | HP | 0.76 | 0.007 | 0.033 | 0.55 | <0.001 | <0.001 | 0.104 | -0.743 |
| HPT (241) - *5402* | HPT | HP | 0.8 | <0.001 | <0.001 | 0.75 | <0.001 | <0.001 | -0.907 | -0.131 |
| HPT (241) - *5511* | HPT | HP | 0.88 | 0.005 | 0.027 | 0.85 | 0.002 | 0.008 | -0.318 | -0.249 |
| HPT (241) - *6502* | HPT | HP | 1.25 | 0.004 | 0.021 | 1.68 | <0.001 | <0.001 | -0.417 | 0.508 |
| IGA2 (205) - 5510 | IGA2 | IGHA2 | 0.42 | 0.003 | 0.016 | 0.08 | <0.001 | <0.001 | -0.592 | -0.558 |
| IGG2 (297) - *4400* | IGG2 | IGHG2 | 0.69 | 0.004 | 0.022 | 0.52 | <0.001 | <0.001 | -0.906 | -0.847 |
| IGG2 (297) - *4411* | IGG2 | IGHG2 | 1.24 | 0.008 | 0.035 | 1.59 | <0.001 | <0.001 | 0.371 | 0.181 |
| IGM (209) - *5401* | IGM | IGHM | 1.5 | 0.003 | 0.016 | 1.47 | 0.011 | 0.031 | 0.17 | 0.266 |

| | | | | | | | | | | |
|---|---|---|---|---|---|---|---|---|---|---|
| KLKB1 (494) - *5401* | KLKB1 | KLKB 1 | 1.7 | <0.001 | 0.001 | 2.87 | <0.001 | <0.001 | 0.017 | 0.086 |
| KLKB1 (494) - *5402* | KLKB1 | KLKB 1 | 1.84 | 0.003 | 0.017 | 2.99 | <0.001 | <0.001 | 0.223 | -0.331 |
| KLKB1 (494) - *5410* | KLKB1 | KLKB 1 | 1.27 | 0.01 | 0.041 | 1.79 | <0.001 | <0.001 | -0.037 | 0.829 |
| KLKB1 (494) - *6503* | KLKB1 | KLKB 1 | 1.51 | <0.001 | 0.002 | 1.6 | <0.001 | <0.001 | 0.707 | -0.646 |
| TRFE (432) - *5402* | TRFE | TF | 1.19 | 0.001 | 0.008 | 1.66 | <0.001 | <0.001 | -0.938 | 1.004 |
| TRFE (432) - *6501* | TRFE | TF | 1.24 | <0.001 | 0.001 | 1.47 | <0.001 | <0.001 | 0.204 | 0.146 |
| TRFE (432) - *6502* | TRFE | TF | 1.13 | 0.013 | 0.049 | 1.42 | <0.001 | <0.001 | -0.539 | 0.2 |
| VTNC (169) - *5401* | VTNC | VTN | 0.71 | <0.001 | 0.002 | 0.54 | <0.001 | <0.001 | -0.28 | -0.238 |
| ZA2G (112) - *5402* | ZA2G | AZGP 1 | 1.49 | 0.008 | 0.034 | 2.06 | <0.001 | <0.001 | -0.308 | 0.171 |

**Table S4.**

| Marker | control/NASH (multiplicative difference) | control/ NASH (p-value) | control/ NASH (FDR) | control/HCC (multiplicative difference) | control/HCC (p-value) | control/HCC (FDR) |
|---|---|---|---|---|---|---|
| A1AT_peptide | 1.39 | <0.001 | 0.001 | 1.12 | 0.1 | 0.176 |
| A1AT (107) - *6502* | 0.76 | 0.169 | 0.304 | 0.6 | 0.029 | 0.064 |
| A1AT (107) - *6512* | 1.17 | 0.105 | 0.219 | 2.03 | <0.001 | <0.001 |
| A1AT (107) - *6513* | 1.24 | 0.066 | 0.157 | 1.58 | 0.018 | 0.043 |
| A1AT (271) - *5401* | 0.85 | <0.001 | 0.005 | 0.79 | <0.001 | <0.001 |
| A1AT (271) - *5402* | 0.87 | <0.001 | <0.001 | 0.82 | <0.001 | <0.001 |
| A1AT (271) - *5412* | 0.55 | <0.001 | <0.001 | 0.71 | <0.001 | <0.001 |
| A2MG_peptide | 0.85 | 0.053 | 0.138 | 1.26 | 0.029 | 0.064 |
| A2MG (1424) - *5401* | 0.97 | 0.779 | 0.841 | 0.82 | 0.043 | 0.088 |
| A2MG (1424) - *5402* | 1.07 | 0.385 | 0.542 | 1.58 | <0.001 | <0.001 |
| A2MG (247) - *5200* | 0.91 | 0.26 | 0.411 | 0.65 | <0.001 | <0.001 |
| A2MG (247) - *5402* | 1.24 | <0.001 | 0.002 | 1.47 | <0.001 | <0.001 |
| A2MG (55) - *5402* | 1.17 | 0.003 | 0.016 | 1.34 | <0.001 | <0.001 |
| A2MG (55) - *5411* | 0.96 | 0.586 | 0.712 | 0.69 | <0.001 | <0.001 |
| A2MG (55) - *5412* | 0.94 | 0.391 | 0.545 | 0.67 | <0.001 | <0.001 |
| A2MG (869) - *5200* | 0.95 | 0.313 | 0.474 | 0.74 | <0.001 | <0.001 |
| A2MG (869) - *5401* | 1.05 | 0.366 | 0.531 | 1.13 | 0.045 | 0.092 |
| A2MG (869) - *5402* | 1.13 | 0.023 | 0.071 | 1.17 | 0.015 | 0.037 |

| | | | | | | |
|---|---|---|---|---|---|---|
| A2MG (869) - *6200* | 0.87 | 0.013 | 0.049 | 0.68 | <0.001 | <0.001 |
| A2MG (869) - 6300 | 0.91 | 0.239 | 0.391 | 0.62 | <0.001 | <0.001 |
| AGP1_peptide | 1.13 | 0.22 | 0.367 | 0.8 | 0.019 | 0.046 |
| AGP1 (103) - *6503* | 0.97 | 0.6 | 0.722 | 0.77 | 0.002 | 0.008 |
| AGP1 (33) - *5402* | 0.72 | <0.001 | 0.003 | 0.75 | 0.002 | 0.008 |
| AGP1 (93) - 6502 | 0.82 | 0.009 | 0.038 | 0.69 | <0.001 | <0.001 |
| AGP1 (93) - *6503* | 0.85 | 0.057 | 0.145 | 0.77 | 0.019 | 0.044 |
| AGP1 (93) - *7613* | 1.03 | 0.776 | 0.841 | 1.63 | <0.001 | <0.001 |
| AGP1 (93) - *7604* | 0.83 | 0.022 | 0.069 | 0.5 | <0.001 | <0.001 |
| AGP12 (72) - *6503* | 0.87 | 0.051 | 0.136 | 0.59 | <0.001 | <0.001 |
| AGP12 (72) - *7601* | 1.01 | 0.908 | 0.932 | 1.33 | 0.016 | 0.04 |
| AGP12 (72) - *7613* | 1.01 | 0.943 | 0.955 | 1.55 | <0.001 | 0.003 |
| AGP12 (72) - *7614* | 1.12 | 0.373 | 0.545 | 1.43 | 0.009 | 0.024 |
| CFAH_peptide | 0.89 | 0.028 | 0.084 | 0.9 | 0.067 | 0.127 |
| CFAH (1029) - *5401* | 0.91 | 0.201 | 0.341 | 0.67 | <0.001 | <0.001 |
| CFAH (1029) - *5431* | 0.79 | 0.023 | 0.0708 | 0.65 | <0.001 | <0.001 |
| CFAH (882) - *5411* | 1.2 | 0.095 | 0.203 | 1.49 | <0.001 | <0.001 |
| CFAH (882) - *5401* | 0.83 | 0.058 | 0.146 | 0.56 | <0.001 | <0.001 |
| CFAH (882) - *5402* | 0.95 | 0.048 | 0.129 | 0.88 | 0.006 | 0.017 |
| CFAH (911) - *5421* | 1.29 | 0.015 | 0.053 | 1.35 | 0.009 | 0.024 |
| CFAH (911) - *5401* | 0.86 | 0.082 | 0.183 | 0.61 | <0.001 | <0.001 |
| HPT_peptide | 1.35 | 0.015 | 0.053 | 0.85 | 0.315 | 0.423 |
| HPT (184) - *5401* | 0.834 | 0.162 | 0.295 | 0.48 | <0.001 | <0.001 |
| HPT (184) - *5402* | 1.09 | 0.061 | 0.151 | 1.26 | <0.001 | <0.001 |
| HPT (184) - *5411* | 1.03 | 0.776 | 0.841 | 1.36 | 0.027 | 0.063 |
| HPT (184) - *5412* | 1.14 | 0.176 | 0.312 | 1.78 | <0.001 | <0.001 |
| HPT (184) - *5510* | 0.78 | 0.062 | 0.1501 | 0.47 | <0.001 | <0.001 |
| HPT (207) – *5401+5402* | 0.96 | 0.642 | 0.756 | 0.55 | <0.001 | <0.001 |
| HPT (207) – *5401+6502* | 0.82 | 0.006 | 0.031 | 0.54 | <0.001 | <0.001 |
| HPT (207) – *6502+6503* | 0.72 | 0.005 | 0.025 | 0.56 | <0.001 | <0.001 |
| HPT (207) – *6502+7613* | 0.67 | 0.041 | 0.116 | 0.7 | 0.009 | 0.025 |
| HPT (241) - *5401* | 0.76 | 0.007 | 0.033 | 0.55 | <0.001 | <0.001 |
| HPT (241) - *5402* | 0.8 | <0.001 | <0.001 | 0.75 | <0.001 | <0.001 |
| HPT (241) - *5511* | 0.88 | 0.005 | 0.027 | 0.85 | 0.002 | 0.008 |
| HPT (241) - *6502* | 1.25 | 0.004 | 0.021 | 1.68 | <0.001 | <0.001 |
| HPT (241) - *6512* | 1.44 | 0.014 | 0.051 | 3.63 | <0.001 | <0.001 |
| HPT (241) - *7604* | 1.08 | 0.599 | 0.722 | 1.74 | <0.001 | <0.001 |

**Table S5.**

| Glycoform Code | Glycoform Code (Hex – mannose/galactose, HexNAc – GlcNAc/GalNAc, Fuc – Fucose, NeuAc – Sialylic acid) | Structure |
|---|---|---|
| *1102* | Hex(1) HexNAc(1)Fuc(0)NeuAc(2) | 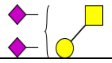 |
| *1202* | Hex(1)HexNAc(2)Fuc(0)NeuAc(2) | 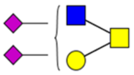 |
| *1300* | Hex(1)HexNAc(3)Fuc(0)NeuAc(0) | 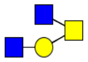 |
| *2110* | Hex(2)HexNAc(1)Fuc(1)NeuAc(0) | 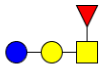 |
| *4400* | Hex(4)HexNAc(4)Fuc(0)NeuAc(0) | 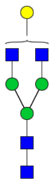 |
| *4411* | Hex(4)HexNAc(4)Fuc(1)NeuAc(1) | 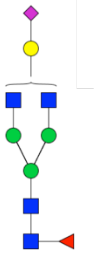 |

| | | |
|---|---|---|
| *5200* | Hex(5)HexNAc(2)Fuc(0)NeuAc(0) | 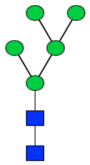 |
| *5400* | Hex(5)HexNAc(4)Fuc(0)NeuAc(0) | 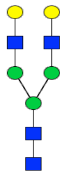 |
| *5401* | Hex(5)HexNAc(4)Fuc(0)NeuAc(1) | 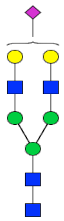 |
| *5402* | Hex(5)HexNAc(4)Fuc(0)NeuAc(2) | 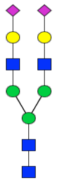 |
| *5410* | Hex(5)HexNAc(4)Fuc(1)NeuAc(0) | 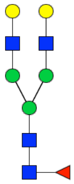 |

| | | |
|---|---|---|
| *5411* | Hex(5)HexNAc(4)Fuc(1)NeuAc(1) | 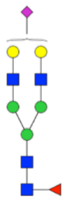 |
| *5412* | Hex(5)HexNAc(4)Fuc(1)NeuAc(2) | 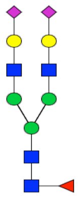 |
| *5421* | Hex(5)HexNAc(4)Fuc(2)NeuAc(1) | 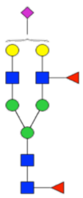 |
| *5431* | Hex(5)HexNAc(4)Fuc(3)NeuAc(1) | 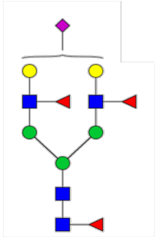 |
| *5510* | Hex(5)HexNAc(5)Fuc(1)NeuAc(0) | 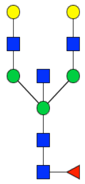 |

| | | |
|---|---|---|
| *5511* | Hex(5)HexNAc(5)Fuc(1)NeuAc(1) | 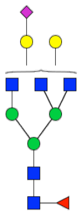 |
| *6200* | Hex(6)HexNAc(2)Fuc(0)NeuAc(0) | 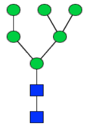 |
| *6300* | Hex(6)HexNAc(3)Fuc(0)NeuAc(0) | 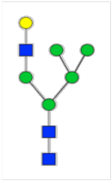 |
| *6500* | Hex(6)HexNAc(5)Fuc(0)NeuAc(0) | 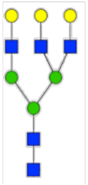 |
| *6501* | Hex(6)HexNAc(5)Fuc(0)NeuAc(1) | 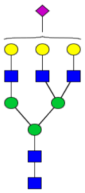 |

| | | |
|---|---|---|
| *6502* | Hex(6)HexNAc(5)Fuc(0)NeuAc(2) | 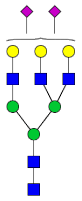 |
| *6503* | Hex(6)HexNAc(5)Fuc(0)NeuAc(3) | 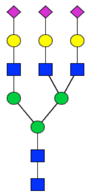 |
| *6511* | Hex(6)HexNAc(5)Fuc(1)NeuAc(1) | 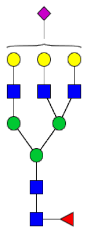 |
| *6512* | Hex(6)HexNAc(5)Fuc(1)NeuAc(2) | 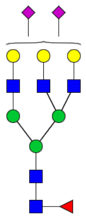 |
| *6513* | Hex(6)HexNAc(5)Fuc(1)NeuAc(3) | 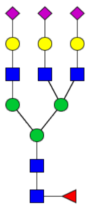 |

| | | |
|---|---|---|
| *7600* | Hex(7)HexNAc(6)Fuc(0)NeuAc(0) | 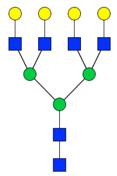 |
| *7601* | Hex(7)HexNAc(6)Fuc(0)NeuAc(1) | 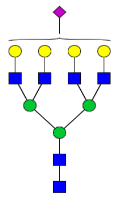 |
| *7601* | Hex(7)HexNAc(6)Fuc(0)NeuAc(2) | 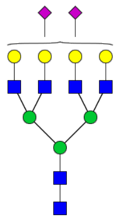 |
| *7604* | Hex(7)HexNAc(6)Fuc(0)NeuAc(4) | 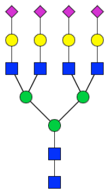 |
| *7613* | Hex(7)HexNAc(6)Fuc(1)NeuAc(3) | 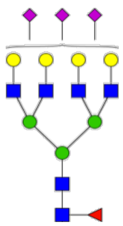 |

| | | |
|---|---|---|
| *7614* | Hex(7)HexNAc(6)Fuc(1)NeuAc(4) | 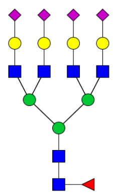 |

**Table S6.**

| Upstream regulator | Molecule type | p-value of overlap | Target molecules in dataset |
|---|---|---|---|
| **SREBF1** | transcription regulator | 7.22E-08 | APOC3, CFAI, IGM, A1AT, AACT, TRFE |
| **IL6** | cytokine | 2.15E-07 | A2MG, CLUS, HPT, IGM, AGP1, A1AT, AACT, TRFE |
| **FOXA2** | transcription regulator | 1.94E-06 | A2MG, APOC3, APOM, A1AT, TRFE |
| **HNF1A** | transcription regulator | 3.18E-06 | APOC3, APOM, CO8A, CFAI, A1AT, VTNC |
| **HNF4A** | transcription regulator | 5.26E-06 | A1BG, APOC3, APOM, CO4A/CO4B, CO6, AGP1, A1AT, AACT, TRFE, VTNC |
| **IL6ST** | transmembrane receptor | 2.32E-05 | A2MG, HPT, AGP1 |
| **PPARA** | ligand-dependent nuclear receptor | 4.91E-05 | APOC3, APOM, CO6, CO8A, CFAI |
| **IL6R** | transmembrane receptor | 5.91E-05 | A2MG, HPT, AACT |
| **JUN** | transcription regulator | 6.27E-05 | A2MG, APOC3, APOM, CLUS, TRFE |
| **IL1A** | cytokine | 6.34E-05 | CO4A/CO4B, AGP1, A1AT, AACT |
